# Spatiotemporal co-dependency between macrophages and exhausted CD8^+^ T cells in cancer

**DOI:** 10.1101/2021.09.27.461866

**Authors:** Kelly Kersten, Kenneth H. Hu, Alexis J. Combes, Bushra Samad, Tory Harwin, Arja Ray, Arjun Arkal Rao, En Cai, Kyle Marchuk, Jordan Artichoker, Tristan Courau, Quanming Shi, Julia Belk, Ansuman T. Satpathy, Matthew F. Krummel

## Abstract

T cell exhaustion is a major impediment to anti-tumor immunity. However, it remains elusive how other immune cells in the tumor microenvironment (TME) contribute to this dysfunctional state. Here we show that the biology of tumor-associated macrophages (TAM) and exhausted T cells (T_ex_) in the TME is extensively linked. We demonstrate that *in vivo* depletion of TAM reduces exhaustion programs in tumor-infiltrating CD8^+^ T cells and reinvigorates their effector potential. Reciprocally, transcriptional and epigenetic profiling reveals that T_ex_ express factors that actively recruit monocytes to the TME and shape their differentiation. Using lattice light sheet microscopy, we show that TAM and CD8^+^ T cells engage in unique long-lasting antigen-specific synaptic interactions that fail to activate T cells, but prime them for exhaustion, which is then accelerated in hypoxic conditions. Spatially resolved sequencing supports a spatiotemporal self-enforcing positive feedback circuit that is aligned to protect rather than destroy a tumor.

**Graphical Abstract:** 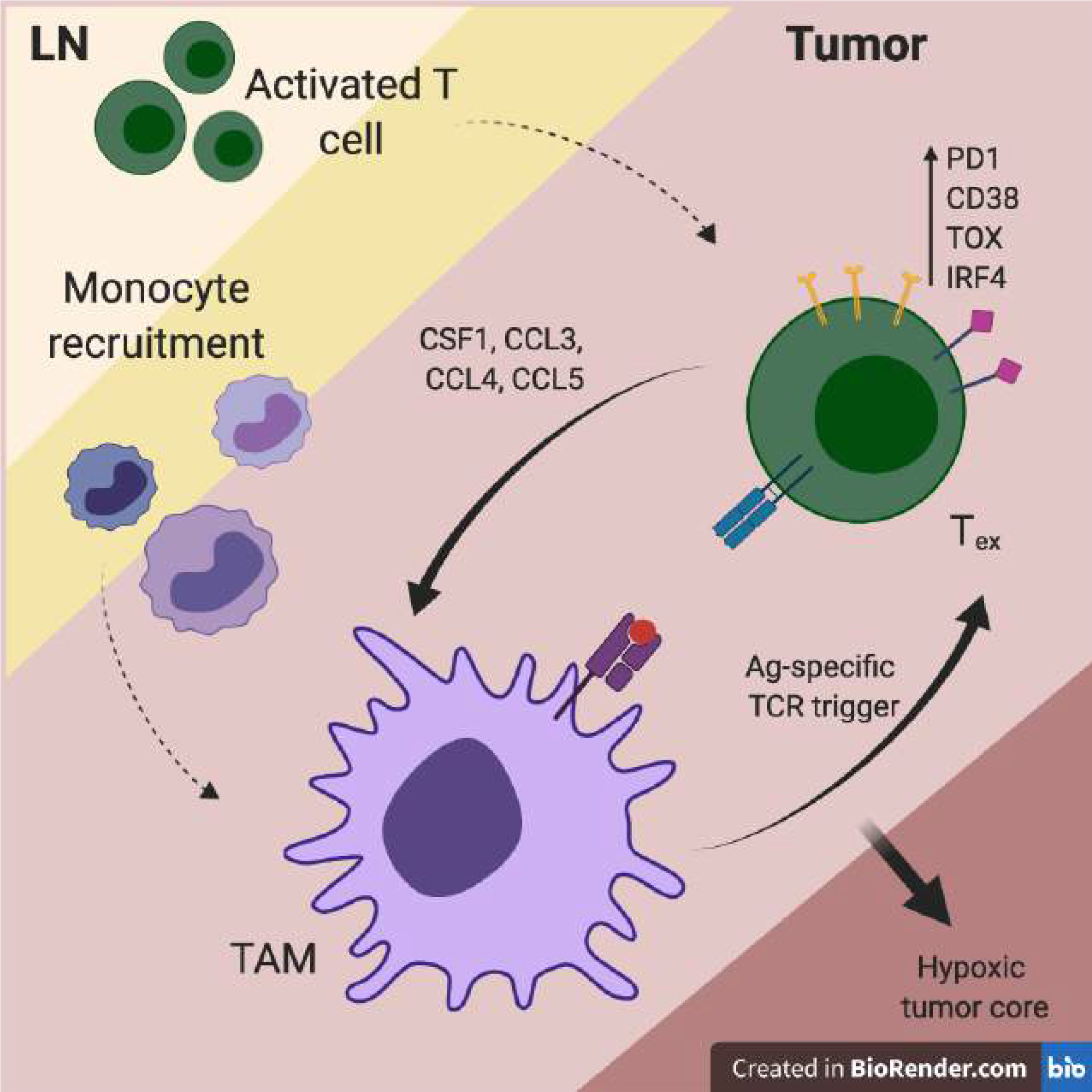

## Introduction

Cancer immunotherapy – harnessing the patient’s immune system to fight cancer – has revolutionized cancer treatment strategies. However, a large proportion of patients does not show clinical response, and the mechanisms underlying intrinsic and acquired resistance are still poorly understood. CD8^+^ T cells are critical mediators of anti-tumor immune responses, and the main target for current immunotherapy approaches. Tumor infiltration of CD8^+^ T cells correlates with improved prognosis and beneficial responses to immune checkpoint blockade as compared to non-infiltrated tumors (Galon et al., 2006; Tumeh et al., 2014). However, those CD8^+^ T cells are frequently non-functional due to establishment of a state of exhaustion, characterized by the expression of inhibitory molecules including PD1, CD38 and TOX and the loss of cytotoxic effector function (Wherry et al., 2007; Doering et al., 2012; Schietinger et al., 2016; Pauken et al., 2016; Scott et al., 2019; Khan et al., 2019). Several studies have shown that chronic antigen exposure and stimulation of the T cell receptor (TCR) are required for exhaustion programs in T cells (Utzschneider et al., 2016; Scott et al., 2019; Oliveira et al., 2021). However, how this is orchestrated in the tumor microenvironment (TME) is unclear.

The immune composition of the TME plays an important role in regulating effective anti-tumor T cell responses (Binnewies et al., 2018). Across solid tumors, the majority of immune cells in the TME is frequently comprised of antigen-presenting myeloid cells (APC), of which tumor-associated macrophages (TAM) are typically the most abundant (DeNardo et al., 2011; Ruffell et al., 2012; Broz et al., 2014). TAM abundance is correlated with poor prognosis in a variety of solid tumor types (Zhang et al., 2012; Gentles et al., 2015), and many studies report on their immunosuppressive role in cancer progression and dissemination (DeNardo and Ruffell, 2019). Conversely, some studies report immunostimulatory and anti-tumor functions of TAM through expression of TNF and iNOS, or upon treatment with CD40 agonists (Beatty et al., 2011; Klug et al., 2013). Similar to a rare population of dendritic cells (cDC1), which have been described to be potent activators of anti-tumor T cells (Broz et al., 2014; Roberts et al., 2016; Salmon et al., 2016; Spranger et al., 2017), TAMs have the potential to phagocytose large amounts of tumor-associated antigens, but fail to successfully support T cell activation (Engelhardt et al., 2012; Broz et al., 2014). Interestingly, intravital imaging studies have shown that antigen-specific CD8^+^ T cells preferentially localize in TAM-rich areas in the TME, and form tight interactions that persist over time (Boissonnas et al., 2013; Broz et al., 2014; Peranzoni et al., 2018).

Here we dissect the molecular mechanisms of a novel immune co-differentiation, by which TAM and exhausted CD8^+^ T cells (T_ex_) sustain each other’s maturation and presence in the TME through long-lived antigen-specific synaptic contacts. Our study reveals a novel mechanistic link through an antigen-driven positive feedback loop and offers a possible path whereby T cells and TAM might equally contribute to initial and sustained tumor immune evasion.

## Results

### CD8^+^ T cell exhaustion correlates with macrophage abundance in the TME

To study how myeloid immune cells contribute to CD8^+^ T cell exhaustion in the TME, we first focused on the concurrent events during the onset of exhaustion programs in tumor-infiltrating CD8^+^ T cells in mouse models of melanoma (B78ChOVA and B16ChOVA) and spontaneous breast cancer (*MMTV-PyMTChOVA*) (Fig. S1A). At different time points during tumor growth, we adoptively transferred Ovalbumin (OVA)-specific OT-I CD8^+^ T cells into tumor-bearing mice, focusing on: (1) early arrival T cells that were recently recruited to the TME (T_ex_ d4) and (2) T cells that have resided in the TME for 14 days and have demonstrably upregulated PD1, CD38, TOX and CD5 as assessed by flow cytometry (T_ex_ d14) (Fig. S1B). Both of these populations demonstrated reduced production of the cytokines IFN*γ* and TNF*α* when compared to activated CD44^+^ endogenous CD8^+^ T cells in the tumor-draining lymph node (TdLN) (Fig. S1C, D). The onset of exhaustion and dysfunction in CD8^+^ T cells upon tumor infiltration was antigen-specific, because irrelevant LCMV-specific P14 CD8^+^ T cells did not acquire phenotypic markers of exhaustion when compared to OT-I CD8^+^ T cells (Fig. S1E-H). However, the ability to produce effector cytokines IFN*γ* and TNF*α* was blunted equivalently in endogenous, P14 and OT-I CD8^+^ T cells upon tumor residence (Fig. S1I, J). This may occur if loss of those markers represented a natural decay process post-activation or if additional non-antigen-specific and more universal suppression mechanisms are at work in the TME.

Tumor-associated macrophages (TAM) comprised the majority of myeloid cells in B78ChOVA melanomas in line with previous studies (Broz et al., 2014; Gentles et al., 2015; Cheng et al., 2021), and their abundance increased during tumor progression, while the fraction of CD103^+^ cDC1 and CD11b^+^ cDC2 diminished (Fig. S1K). Recognizing the stoichiometric abundance of TAM as possible APC, we sought to study the role of TAM in the onset of CD8^+^ T cell exhaustion by subjecting tumor-bearing mice to antibody-mediated blockade of CSF1/CSF1R signaling (Fig. 1A and Fig. S1L, O). This treatment resulted in significant reduction of the proportion of CD11b^+^F4/80^+^ TAM in tumors (Fig. 1B,C and Fig. S1M, P). TAM-depletion also resulted in a concurrent reduction in the expression of exhaustion markers PD1, CD38 and TOX on tumor-infiltrating CD44^+^ OT-I CD8^+^ T cells (Fig. 1D and Fig. S1N, Q). Moreover, the resultant CD44^+^ OT-I CD8^+^ T cells produced higher levels of IFN*γ* and TNF*α* in anti-CSF1R-treated compared to isotype-treated mice (Fig. 1E, F). We also found that the expression of PD1, CD38 and TOX on tumor-infiltrating CD44^+^ OT-I CD8^+^ T cells positively correlates with the abundance of TAM in the TME, measured across 25 mice in three independent experiments subjected to anti-CSF1R or isotype treatment (Fig. 1G). In line with this, flow cytometric profiling of CD8^+^ tumor-infiltrating lymphocytes (TIL) in a cohort of 20 patients with renal cell carcinomas – which are rich in myeloid and T cells (Combes et al., 2021; Mujal et al., 2021) –, demonstrated a strong association between PD1 and CD38 (but not CTLA4) expression and the degree to which patient’s myeloid cells had differentiated toward macrophages as compared to monocytes in the TME (Fig. 1H). We used this ratio as the pure number of macrophages did not show this association (data not shown).

**Figure 1.**
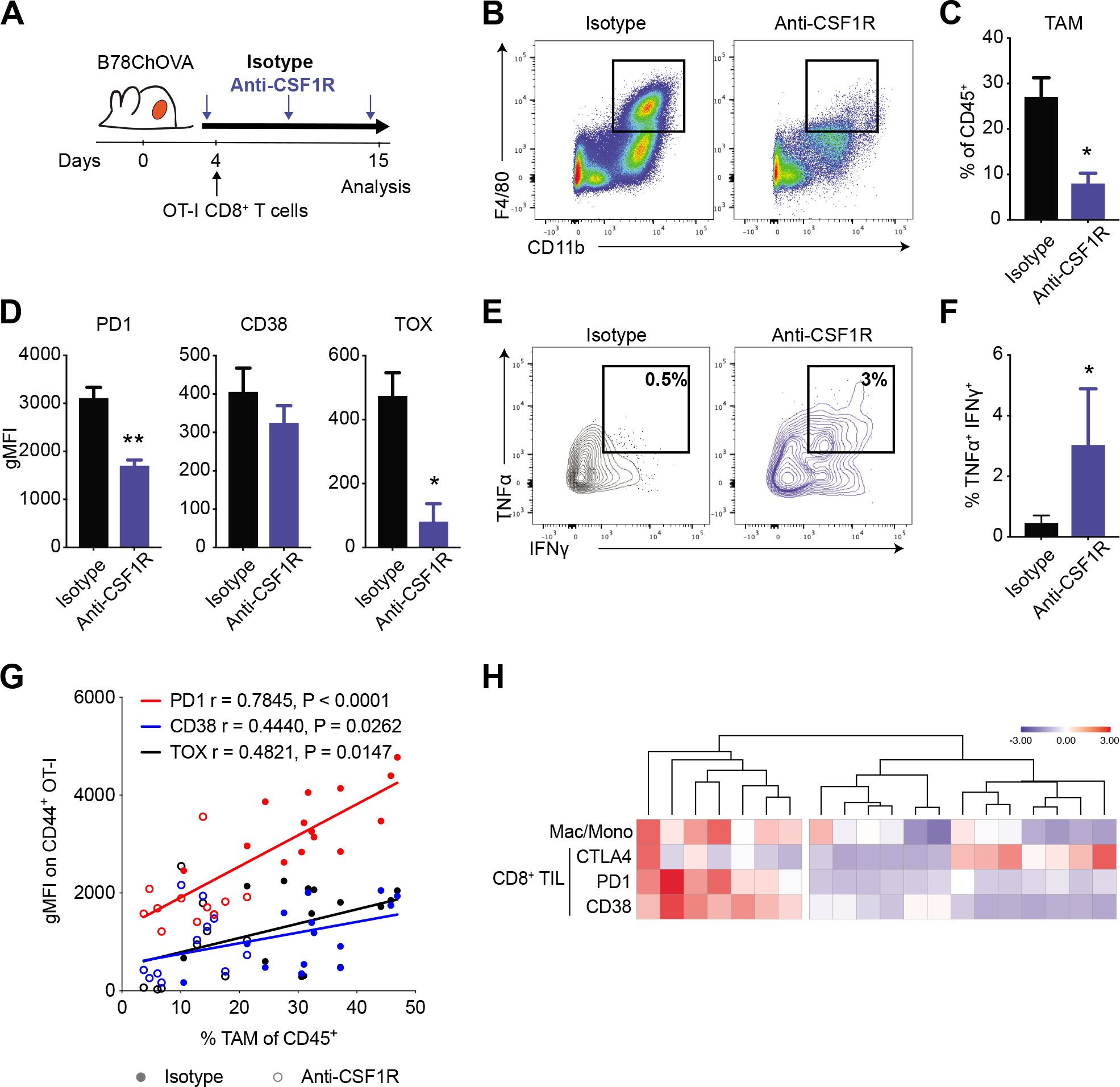
CD8^+^ T cell exhaustion correlates with macrophage abundance in the TME. A) Experimental setup to study the role of macrophages in CD8^+^ T cell exhaustion in B78ChOVA melanoma. Weekly anti-CSF1R or isotype antibody treatment was initiated 1-2 days after tumor inoculation and continued until mice were sacrificed. OVA-specific OT-I CD8^+^ T cells were adoptively transferred 2-3 days after tumor inoculation. Mice were sacrificed at day 15 and tumors were harvested for analysis. B-C) Representative flow plots (B) and quantification (C) of CD11b^+^ F4/80^+^ macrophages in isotype and anti-CSF1R-treated B78ChOVA melanomas. N=5 mice/group. D) Surface (PD1 and CD38) and intracellular (TOX) expression on intratumoral CD44^+^ OT-I CD8^+^ T cells from isotype and anti-CSF1R-treated mice. N=5 mice/group. E-F) Representative contour plots (E) and quantification (F) of IFN*γ*^+^TNF*α*^+^ polyfunctional CD44^+^ OT-I CD8^+^ T cells in tumors of isotype and anti-CSF1R-treated mice. N=8-9 mice/group. Pooled data from two independent experiments. G) Spearman correlation between gMFI of PD1, CD38 and TOX expression on CD44^+^ OT-I CD8^+^ T cells and % of TAM of CD45^+^ cells in B78ChOVA melanomas treated with isotype (solid) or anti-CSF1R antibody (open). N=11-14 mice/group. H) Heatmap showing clustering of normalized z-scores of CTLA4, PD1 and CD38 expression on CD8^+^ TIL and macrophage/monocyte ratio in 20 fresh human renal cell carcinoma samples (rows) determined by flow cytometry. All data are mean ± SEM. ** p < 0.01, * p < 0.05 as determined by Mann-Whitney U-test. See also Figure S1.

### CD8^+^ T_ex_ express monocyte/macrophage-related factors upon prolonged residence in the TME of mouse and human cancers

To test whether there might be a mechanistic link between TAM abundance and CD8^+^ T cell exhaustion, we isolated early (T_ex_ d4) and late exhausted OT-I CD8^+^ T cells (T_ex_ d14) from B78ChOVA tumors and compared their transcriptional profile to that of splenic naïve CD44^−^ OT-I CD8^+^ T cells by RNAseq. As expected, T_ex_ d14 showed enhanced expression of known markers associated with exhaustion (Sade-Feldman et al., 2018), including but not limited to *Cd44*, *Pdcd1*, *Cd38*, *Tox*, *Irf4*, *Havcr2*, *Lag3,* while expression of naïve precursor genes *Sell, Tcf7* and *Il7r* were enriched in T_naïve_ cells (Fig. 2A). Interestingly, pathway enrichment analysis of genes with a fold enrichment >5 in T_ex_ revealed dramatic enrichment of pathways involved in “abnormal cytokine secretion” and “impaired macrophage chemotaxis” (Fig. 2B). A closer analysis of individual genes demonstrated that expression of genes associated with naïve precursor T cell states decreased and genes previously associated with exhaustion increased in T_ex_ d14 vs T_ex_ d4 (Fig. 2C), consistent with previous reports (Pauken et al., 2016; Schietinger et al., 2016; Sade-Feldman et al., 2018). Interestingly, a large set of myeloid-related genes was highly upregulated in exhausted CD8^+^ T cells and most of these increased with prolonged residence in the TME (Fig. 2C and Fig. S2A). Increased expression of *Csf1*, *Ccl3* and *Ccl5* transcripts was confirmed by qRT-PCR on an independent sample set of T_naïve_, T_ex_ d4 and T_ex_ d14 (Fig. S2B). Of note, the majority of these transcriptional changes was not observed in *in vitro* activated effector OT-I CD8^+^ T cells (T_eff_) (Fig. 2C), suggesting a unique exhaustion-related transcriptional profile in CD8^+^ T cells that is acquired *in vivo* upon prolonged residence in the TME.

**Figure 2.**
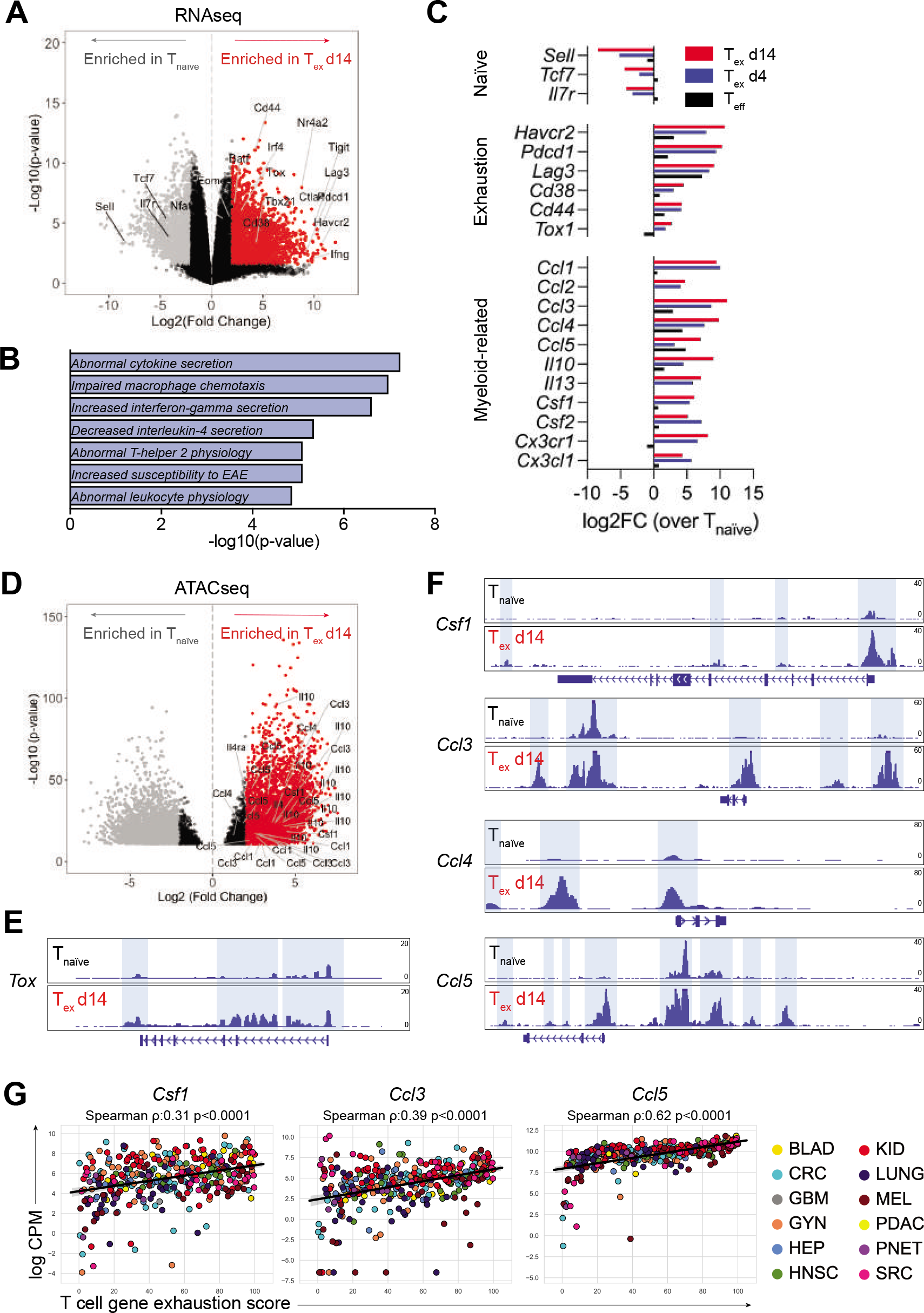
Exhausted CD8^+^ T cells express myeloid-related factors. A) Volcano plot showing differential gene expression in tumor-infiltrating CD44^+^ OT-I CD8^+^ T_ex_ d14 cells (red) compared to splenic CD44^−^ OT-I CD8^+^ T_naïve_ cells (grey) by RNAseq. Colored dots (grey and red) represent genes with a log2FC>2 and FDR<0.05. B) Gene set enrichment analysis of DEGs (log2FC>5 and p-value<0.05) enriched in T_ex_ d14 versus T_naïve_ using the MGI Mammalian Phenotype Level 4 library. C) Average gene expression in T_eff_ (black), T_ex_ d4 (blue) and T_ex_ d14 (red) normalized to T_naïve_ as determined by RNAseq. D) Volcano plot showing differential chromatin accessibility at transcriptional start sites in loci of myeloid genes in tumor-infiltrating CD44^+^ OT-I CD8^+^ T_ex_ d14 cells (red) compared to splenic CD44^−^ OT-I CD8^+^ T_naïve_ cells (grey) by ATACseq. Colored dots (grey and red) represent genes with a log2FC>2 and FDR<0.05. E-F) ATACseq signal tracks at the *Tox* (E), *Csf1*, *Ccl3, Ccl4* and *Ccl5* loci (F) highlighting differential chromatin accessibility peaks in CD44^+^ OT-I CD8^+^ T_ex_ cells (d14) compared to splenic CD44^−^ OT-I CD8^+^ T_naïve_ cells. G) Correlation of normalized expression of *Csf1*, *Ccl3* and *Ccl5* transcripts and exhaustion score in FACS sorted human intratumoral T cells across multiple human cancer indications. See also Figure S2.

Epigenetic profiling using assay for transposase-accessible chromatin with sequencing (ATACseq) confirmed a significant enhancement of overall chromatin accessibility near the transcription start site of the genes encoding these myeloid-related genes in T_ex_ d14 versus T_naïve_ cells (Fig. 2D). A more detailed analysis of signal tracks of chromatin accessibility peaks at different gene loci revealed that *Tox* – a major transcriptional and epigenetic regulator of T cell exhaustion (Alfei et al., 2019; Khan et al., 2019; Scott et al., 2019; Yao et al., 2019) – as well as other well-known genes associated with exhaustion programs (*Pdcd1*, *Cd38*, *Havcr2*, *Ctla4*, *Lag3* and *Entpd1* (Pauken et al., 2016; Philip et al., 2017)) showed increased chromatin accessibility at promoter regions in T_ex_ when compared to T_naïve_ (Fig. 2E and Fig. S2C). This enhanced accessibility was also observed for myeloid-related genes *Csf1*, *Ccl3*, *Ccl4* and *Ccl5* in T_ex_ when compared to T_naïve_ (Fig. 2F). In addition, utilizing transcriptional profiles of T cells isolated from human cancers, we found that increased expression of a T cell exhaustion score correlated significantly with the expression of *Csf1*, *Ccl3* and *Ccl5* in T cells in a dataset comprising hundreds of patients across a dozen cancer indications (Combes et al., 2021) (Fig. 2G).

### CD8^+^ T_ex_ shape the myeloid compartment in mouse melanoma

To directly study the functional significance of chemokine gene expression by T_ex_, we adopted a transwell experimental system using OT-I T cells with varying activation states in the bottom well, and bone marrow-derived monocytes in the upper transwell insert (Fig. 3A). After 24 hrs of culture, a significantly higher number of monocytes had migrated through the transwell membrane towards T_ex_ when compared to T_eff_, T_naïve_ or no T cells (Fig. 3B), demonstrating that T_ex_ actively secrete factors that recruit monocytes while T cells in other activation states do not. Phenotypic analysis by flow cytometry revealed that monocytes co-cultured for 2 days with T_ex_ also show increased uniformity and/or magnitude of expression of CD80, CD86, H2kB and MHCII when compared to T_eff_, T_naïve_ or no T cells (Fig. 3C), suggesting that T_ex_-derived factors augment antigen-presentation potential in differentiating myeloid cells.

**Figure 3.**
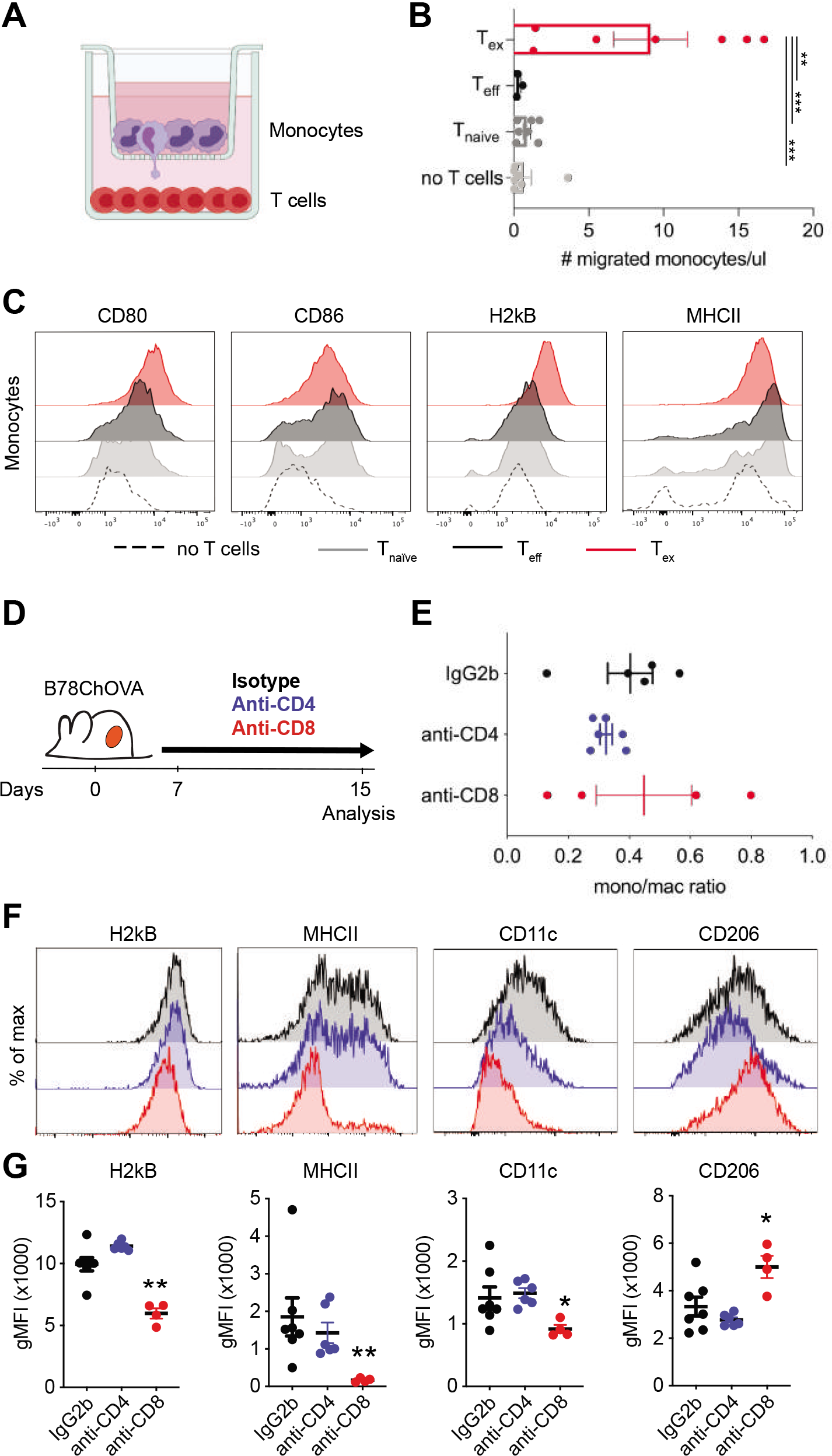
Exhausted CD8^+^ T cells actively recruit monocytes to the TME and shape macrophage phenotype. A) Experimental setup of *in vitro* recruitment assay. Bone marrow-derived monocytes are cultured on transwell inserts (5µm pore size) and T cells (OT-I T_naïve_, T_eff_ and T_ex_) are plated in the bottom well. B) Quantification of recruited monocytes after 24 hours. Data combined from two independent experiments. Statistical significance was determined by one-way ANOVA with Holm-Sidak’s multiple testing correction. C) Representative histograms of expression of surface markers on monocytes after 48 hours of co-culture with T_naïve_, T_eff_ and T_ex_ cells. D) Experimental set-up of *in vivo* CD4^+^ and CD8^+^ T cell depletion in B78ChOVA-bearing mice. Treatment with anti-CD4/CD8 antibodies or isotype was initiated 7 days after tumor inoculation and continued until mice were sacrificed. E-F) Monocyte/macrophage ratio of the proportion of Ly6C^hi^ monocytes and F4/80^+^ macrophages (gated of CD45^+^ cells) in the B78ChOVA TME after isotype, anti-CD4 and anti-CD8 treatment. F-G) Representative histograms (F) and quantification (G) of H2kb, MHCII, CD11c and CD206 expression on CD11b^+^F4/80^+^ macrophages in B78ChOVA tumors after isotype, anti-CD4 and anti-CD8 treatment. Statistical significance was determined using the Mann-Whitney U test. All data are mean ± SEM. * p < 0.05, ** p < 0.01, *** p < 0.001. See also Figure S3.

To assess whether T_ex_ actively shape the myeloid compartment in tumors *in vivo*, we were faced with a dearth of available models that allow for specific (conditional) deletion of exhausted T cells from the TME. Therefore, we decided to use a more widely used approach to systemically deplete CD4^+^ or CD8^+^ T cells from B78ChOVA-bearing mice using depleting antibodies (Fig. 3D). Interestingly, while the ratio of monocytes to macrophages did not significantly change upon CD4- or CD8-depletion (Fig. 3E), their phenotype was drastically affected by CD8^+^ T cell depletion in ways consistent with the results of our *in vitro* co-cultures. CD11b^+^F4/80^+^ TAM showed significantly reduced expression of H2kB, MHCII and CD11c in the absence of CD8^+^ T cells, but not CD4^+^ T cells (Fig. 3F, G). In addition, CD8^+^ T cell depletion resulted in increased expression of ‘pro-tumorigenic M2-marker’ CD206 on TAM (Fig. 3F, G), suggesting that CD8^+^ T cells specifically shape myeloid cell phenotype in the TME favoring an ‘M1-like’ antigen-presenting state.

To test whether the production of CSF1 by lymphocytes had any functional relevance for myeloid composition in the TME, we generated mixed bone marrow chimeras in which *Rag1^−/−^* bone marrow was mixed 50:50 with either *CSF1^op/op^* or *CSF1^op/+^* bone marrow and transferred into lethally irradiated *Rag1^−/−^* recipient mice (Fig. S3A). After a recovery period of 6-10 weeks, these mice were inoculated with B78ChOVA melanomas for 21 days after which the myeloid compartment in the TME was analyzed by flow cytometry. CSF1-deficiency in lymphocytes (*Rag1^−/−^:CSF1^op/op^* chimeras) did not affect primary tumor growth (Fig. S3B) and modestly reduced the influx of total CD45^+^ leukocytes compared to control animals (*Rag1^−/−^:CSF1^op/+^*) (Fig. S3C). In the myeloid compartment, the proportion of Ly6C^+^ monocytes was significantly enriched in *Rag1^−/−^:CSF1^op/op^* chimeras, while macrophage proportions were lower, resulting in an increased monocyte/macrophage ratio in *Rag1^−/−^:CSF1^op/op^* chimeras (Fig. S3D). Of note, the proportion of CD103^+^ cDC1 and CD11b^+^ cDC2 were not appreciably modulated by CSF1-deficiency in lymphocytes (Fig. S3E). In line with the results presented in Fig. 3F and G, the levels of H2kB, MHCII and CD11c on CD11b^+^F4/80^+^ TAM were lower in *Rag1^−/−^:CSF1^op/op^* chimeras when compared to *Rag1^−/−^:CSF1^op/+^* chimeras (Fig. S3F), while expression of CD86 and CD206 were modestly increased. Despite the large biological variation among these samples, the results support the notion that exhausted CD8^+^ T cells, at least partially through expression of CSF1, contribute to TAM maturation and induce an antigen-presentation phenotype in the TME.

### Macrophages and CD8^+^ T cells engage in unique long-lived interactions and synapse formation

Since our data suggests that CD8^+^ T_ex_ shape myeloid cell phenotype towards an antigen-presenting state, we took a more detailed look at the interactions between TAM and CD8^+^ T cells in the TME. Using 2-photon microscopy, we have previously shown that newly infiltrated antigen-specific CD8^+^ T cells in the TME preferentially localize in TAM-rich areas, and are captured in prolonged interactions with TAM that result in the onset of exhaustion programs (Engelhardt et al., 2012; Broz et al., 2014; Boldajipour et al., 2016). Conventional wide field imaging demonstrated that, also *ex vivo* and outside of the context of the TME, previously activated OT-I CD8^+^ T cells interact significantly longer with TAM sorted from OVA-expressing tumors as compared to *in vitro* generated bone marrow-derived dendritic cells (BMDC) unloaded or loaded with the cognate peptide SIINFEKL (SL8) (Fig. 4A, B), suggesting a stable and persistent interaction between TAM and CD8^+^ T cells despite their consistent inability to stimulate T cell proliferation.

**Figure 4.**
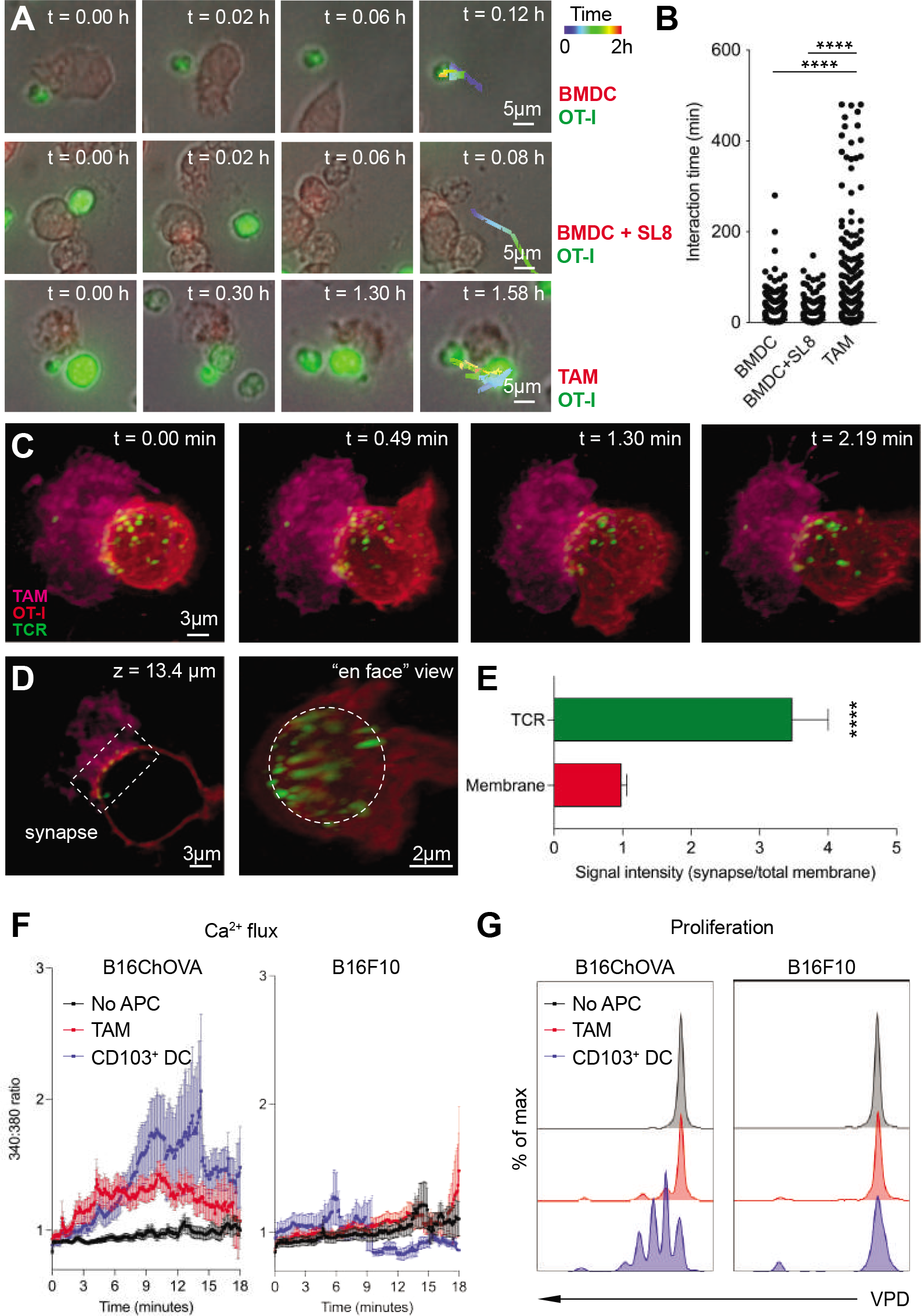
TAM uniquely engage CD8^+^ T cells in antigen-specific long-lived synaptic interactions. A) Representative images of *ex vivo* interactions between mTomato^+^ APC (BMDC, BMDC+SL8 or B78ChOVA-derived TAM) and CFSE-labeled previously activated CD8^+^ OT-I T cells over time using conventional wide field microscopy. B) Quantification of interaction time. n = 3452 TAM, n = 6134 BMDC, n = 3320 BMDC+SL8. Statistical significance was determined using the one-way ANOVA test with Holm-Sidak’s multiple comparison correction. C) Representative images of the interaction between mTomato^+^ TAM sorted from B78ChOVA melanomas (magenta) and previously activated CD8^+^ OT-I T cell labeled with CD45-AF647 (red), with the H57 TCR*β* labeled with AF488 by lattice light sheet imaging. Scale bar, 3µm. D) Z-slice (left) and ‘en face’ view (right) of the TAM-CD8^+^ T cell interaction site showing TCR clustering in the immunological synapse (box (left) and dotted circle (right)). E) Quantification of polarized TCR clustering by determining the ratio of signal intensity of the red (membrane) or green channel (TCR) at the synapse site normalized to the entire membrane. N = 12 T cells. Statistical significance was determined using the Mann-Whitney U test. F-G) Quantification of immediate Ca^2+^ flux by FURA-2AM imaging (F) and proliferation after 72 hours by dilution of Violet Proliferation Dye (VPD) (G) in previously activated CD8^+^ OT-I T cells after interaction with TAM or CD103^+^ DC isolated from B16ChOVA or B16F10 tumors. Negative control represents CD8^+^ OT-I T cells that did not touch an APC (no APC). All data are mean ± SEM. * p < 0.05, ** p < 0.01, *** p < 0.001, **** p < 0.0001. See also Figure S4.

Using lattice light sheet microscopy, we found that this stable interaction between TAMs and CD8^+^ T cells results in small-scale clustering of TCR at the TAM-interaction site on CD8^+^ T cells (Fig. 4C-E and Fig. S4A), consistent with this being a signaling interaction. Calcium imaging revealed that, unlike CD103^+^ cDC1 – potent inducers of CD8^+^ T cell activation (Broz et al., 2014; Roberts et al., 2016; Salmon et al., 2016; Spranger et al., 2017) – that trigger a transient flux, B16ChOVA-derived TAM induce a weak, but long-lasting Ca^2+^ flux in CD8^+^ T cells upon recognition of cognate antigen (Fig. 4F (left)). This flux was likely antigen-dependent since TAM and CD103^+^ cDC1 isolated from non-OVA-expressing B16F10 melanomas did not induce a TCR trigger in OT-I CD8^+^ T cells (Fig. 4F (right)). While the transient Ca^2+^ flux triggered by CD103^+^ cDC1 is sufficient to induce proliferation of CD8^+^ T cells, the antigen-specific trigger provided by TAM fails to support proliferation (Fig. 4G). Thus, despite actively and profoundly engaging T cells in a unique long-lasting antigen-specific signal-generating synaptic interaction, TAM fail to fully support CD8^+^ T cell activation and proliferation.

### Unique TCR engagement by TAM induces exhaustion programs in CD8^+^ T cells

To take a more detailed look at the TAM-induced ‘dysfunctional’ TCR trigger, we examined *ex vivo* co-cultures of previously activated OT-I CD8^+^ T cells with TAM or CD103^+^ cDC1 isolated from B16ChOVA and B16F10 melanomas, or *in vitro* generated BMDC devoid of any antigen as a negative control. As expected, after 3 days of co-culture only CD8^+^ T cells that had encountered CD103^+^ cDC1 expressing their cognate antigen displayed a CD44^hi^ IRF4^hi^ fully activated phenotype (Fig. 5A). Interestingly, when compared to the successful signal provided by CD103^+^ cDC1, TAM only modestly induced expression of activation marker CD44, while providing a similar strength in TCR trigger as suggested by the level of IRF4 expression (Fig. 5A). In chronic viral infections, IRF4 has been implicated as a main regulator of transcriptional circuits inducing and sustaining T cell exhaustion (Man et al., 2017). In line with this notion, TAM induced a significant increase in expression of PD1 and TOX in CD8^+^ T cells in an antigen-specific manner and similar to CD103^+^ cDC1 (Fig. 5A), but nevertheless fail to support proliferation (Fig. 4G).

**Figure 5.**
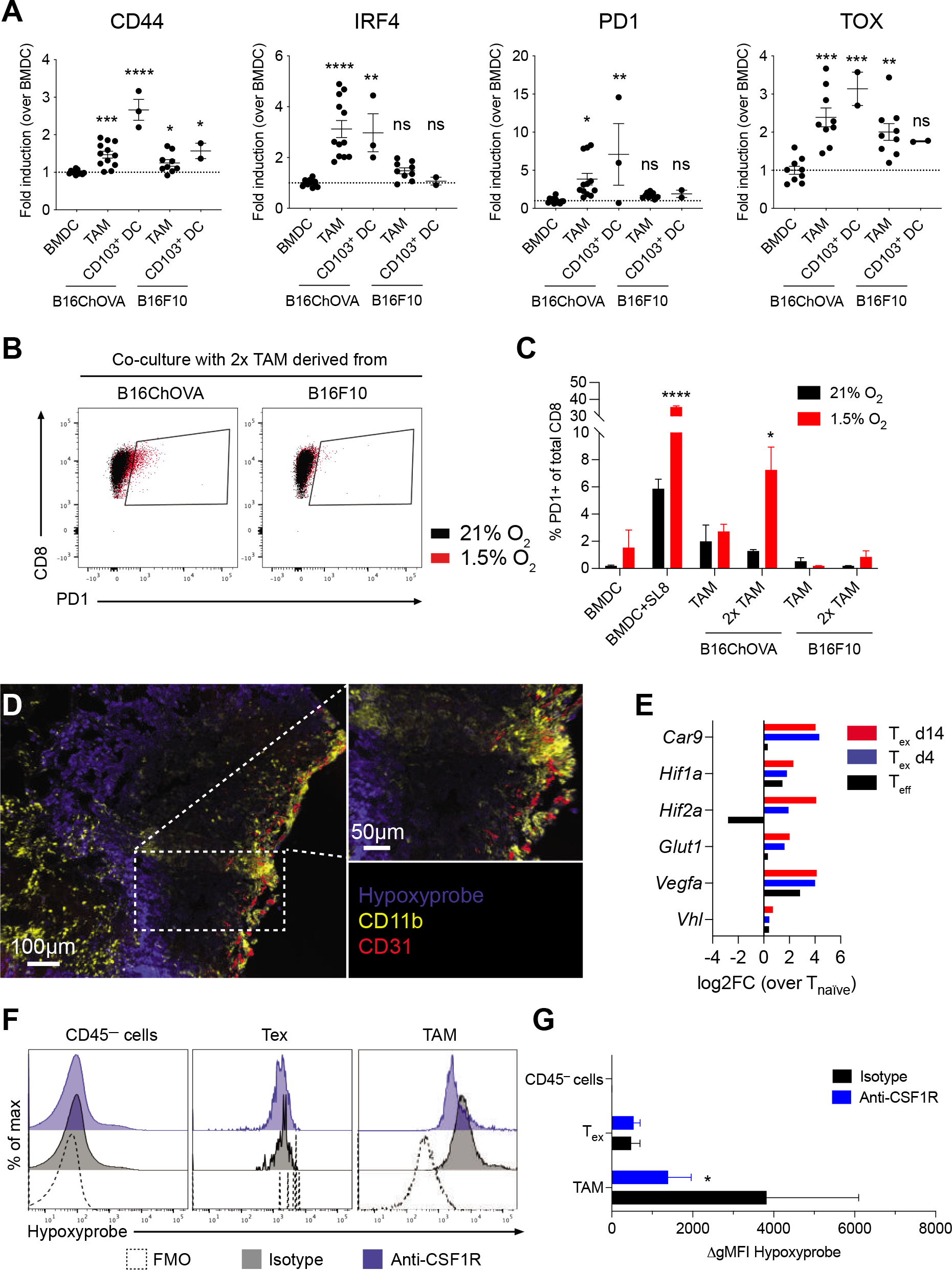
TAM engagement contributes to induction of exhaustion programs in CD8^+^ T cells in an antigen-specific manner. A) Flow cytometric analysis of CD44, IRF4, PD1 and TOX expression in previously activated CD8^+^ OT-I T cells co-cultured for 72 hours with *in vitro* generated BMDC, and TAM or CD103^+^ DC isolated from B16ChOVA or B16F10 tumors. Data presented as fold induction over BMDC. Cumulative data from 4 independent experiments. All data are plotted as mean ± S.E.M. One-way ANOVA with Holm-Sidak correction for multiple comparisons. B-C) Representative dot plots (B) and quantification (C) of PD1^+^ expression on previously activated CD8^+^ OT-I T cells after co-culture with *in vitro* generated BMDC±SL8, and TAM isolated from B16ChOVA or B16F10 tumors. Ratio of APC:T cell was 1:4 or 1:2 (in case of 2x TAM). Plates were incubated in normoxic (21% O_2_) and hypoxic (1.5% O_2_) conditions for 3 days prior to flow cytometric analysis. Statistical significance was determined using the Unpaired t-test. D) Immunofluorescence of B78ChOVA melanomas stained with pimonidazole (Hypoxyprobe) Pacific Blue, CD11b-AF594 (yellow) and CD31-AF647 (red). E) Average expression of hypoxia-related genes in T_eff_ (black), T_ex_ d4 (blue) and T_ex_ d14 (red) normalized to T_naïve_ as determined by RNAseq. F-G) Representative histograms (F) and quantification (G) of hypoxyprobe staining in CD45^—^ cells, CD44^+^ OT-I CD8^+^ T cells and CD11b^+^F4/80^+^ TAM in B78ChOVA melanomas treated with isotype (black) or anti-CSF1R antibodies (blue). Statistical significance was determined using the Mann-Whitney U test. N = 4-5 mice/group. Representative of two independent experiments. All data are mean ± SEM. * p < 0.05, ** p < 0.01, *** p < 0.001, **** p < 0.0001.

Previous work has shown that hypoxia – in combination with chronic antigenic stimulation but not in the absence of stimulation – is required to obtain a ‘full blown’ exhausted phenotype in CD8^+^ T cells *in vitro* (Scharping et al., 2021). To study whether the signals from TAM-T_ex_ interactions are sufficient to ‘prime’ T cells for exhaustion despite being deficient at inducing their proliferation, we performed co-culture experiments under hypoxic (1.5% O_2_) and normoxic (21% O_2_) conditions. Interestingly, the proportion of PD1^+^ CD8^+^ T cells induced by B16ChOVA-derived TAM was much more pronounced when cultured in hypoxic conditions when compared to normoxia, especially when TAM were numerically in excess (Fig. 5B, C). Together these data demonstrate that TAM can prime the onset of exhaustion programs in CD8^+^ T cells, a process that is exacerbated in hypoxic conditions.

Using the hypoxia tracer pimonidazole, we found that B78ChOVA melanomas show highly hypoxic areas towards the inner regions of the tumor, and away from CD31^+^ blood vessels (Fig. 5D). Moreover, we find that these hypoxic regions are surrounded by patches of CD11b^+^ macrophages (Fig. 5D). In line with this observation, expression of hypoxia-related genes (*Car9, Hif1a, Hif2a, Glut1, Vegfa, Vhl*) is upregulated in exhausted CD8^+^ T cells upon prolonged residence in the TME (Fig. 5E), suggesting that tumor-infiltrated T_ex_ experience severe hypoxia in the TME. Interestingly, flow cytometric analysis revealed that TAM experience more severe levels of hypoxia compared to exhausted CD8^+^ T cells and CD45^—^ tumor cells (Fig. 5F, G). Moreover, the degree of hypoxia was dramatically reduced in residual TAM, but not in T_ex_, after CSF1R blockade (Fig. 5F, G).

### ZipSeq mapping reveals spatial coordination of TAM-T_ex_ interaction dynamics in the TME

To better understand the spatial coordination of the dynamic interplay between TAM and T_ex_ in the TME, we utilized ZipSeq, a spatial transcriptomics approach that allows us to map gene expression patterns in single cells based on their localization in the TME by printing barcodes directly onto cells in tissue (Hu et al., 2020). We used a novel *CD206-LSL-Venus-DTR* mouse model in which expression of a Venus fluorescent reporter is driven by the endogenous *CD206* promoter (Fig. S5A). When crossed to the CSF1R^Cre^ strain, we found Venus-labeling of the majority of tumor-associated myeloid cells in B78ChOVA melanomas (Ray A. *et al*, in preparation). We utilized this model to clearly define distinct regions in the tumor such as the outer rim, middle and inner compartment based on the mCherry signal in cancer cells and the Venus-expression in CD206^+^ myeloid cells (Fig. 6A) and apply unique Zipcodes to the surface of immune cells in each of those regions. After dissociation of tumors, we sorted CD45^+^ immune cells and encapsulated them for our modified 10x Genomics scRNA-seq workflow (Hu et al., 2020).

**Figure 6.**
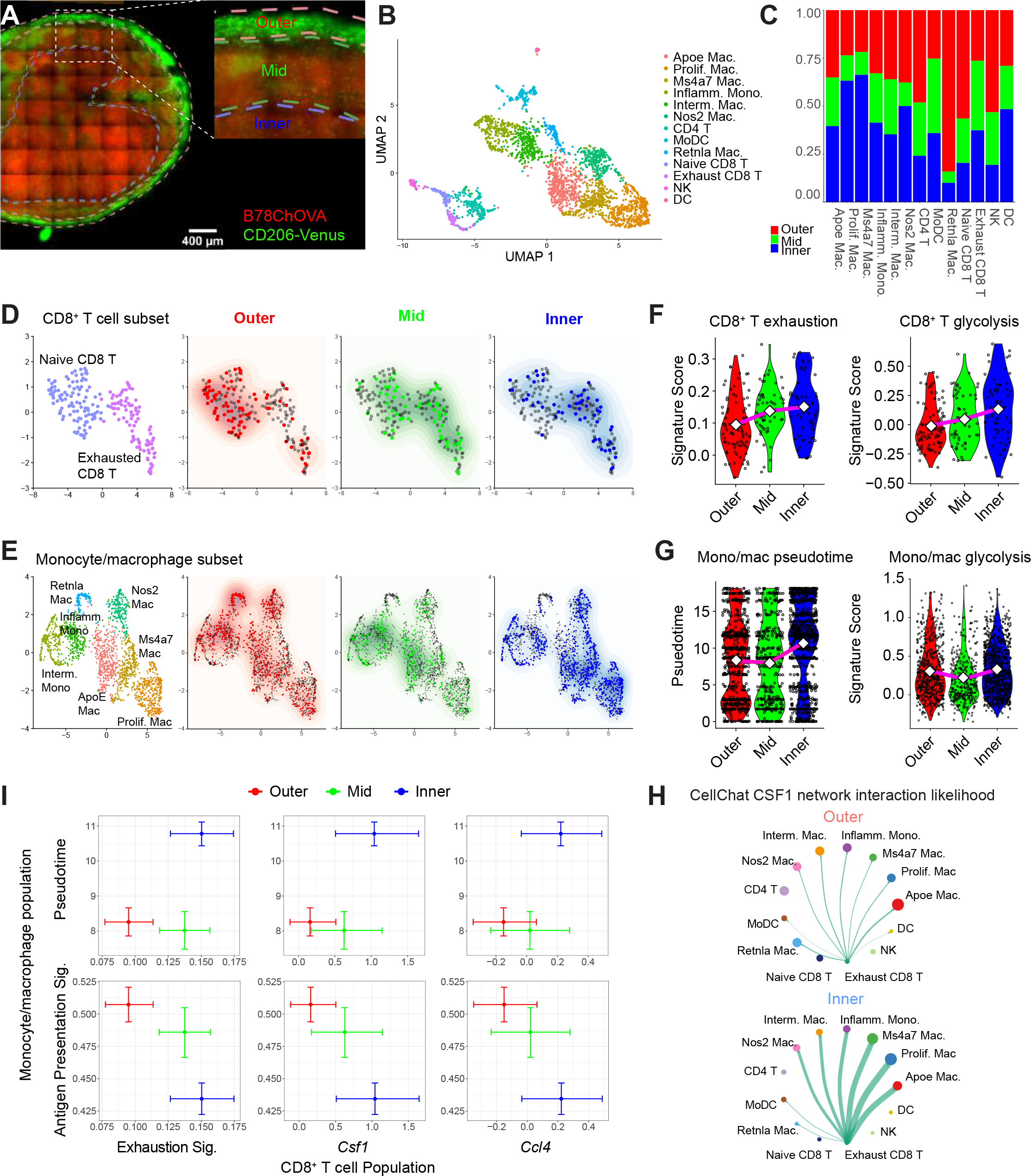
Spatial delineation of TAM-T_ex_ interaction dynamics in the TME. A) Imaging of a 150um-thick live B78ChOVA melanoma section with ROI demarcation of outer, mid and inner compartments used for subsequent ZipSeq. Red channel denotes mCherry signal from B78ChOVA cancer cells and green channel indicates expression of mVenus in CD206^+^ macrophages in the *CSF1R^Cre^;LSL-CD206-Venus-DTR* mouse model. Scale bar = 400µm. B) UMAP representation of sorted CD45^+^ cells following 10X Genomics scRNAseq workflow (n = 2765 cells with n = 427/394/335/288/275/244/220/170/120/108/91/62/31 for clusters as listed). C) Stacked bar charts representing regional distribution of distinct populations identified in B. D-E) UMAP of subsampled CD8^+^ T cell subset (n = 199 cells) (D) and monocyte/macrophage subset (n = 2083 cells) (E) overlaid with their regional localization (Outer, Mid and Inner). F) Violin plots representing CD8^+^ T cell exhaustion score (left) (n = 199 cells) and CD8^+^ T cell glycolytic score (right) (n = 199) in distinct regions in the TME. G) Violin plots representing monocyte/macrophage pseudotime signature score (left) (n = 2083 cells) and monocyte/macrophage glycolytic score (right). H) CellChat interaction likelihood analysis for CSF1 network in outer (top) and inner (bottom) regions of the TME. Thickness of green arrows represents interaction likelihood between populations. I) Cross-whisker plots comparing expression of exhaustion signature and normalized single gene (*Csf1* and *Ccl4)* expression in CD8^+^ T cells (x-axis), and pseudotime score and antigen-presentation signature in the monocyte/macrophage population (y-axis) in distinct regions in the TME. Error bars represent 95% CI as computed by bootstrap resampling. See also Figure S5 and S6 and Table S1.

uMAP analysis of the entire immune compartment revealed prototypical and predominant clusters of T cells and monocytes/macrophages, and smaller clusters of natural killer (NK) cells and dendritic cells (DC) (Fig. 6B and Fig. S6A, B). Some of these populations were enriched in specific regions in the TME (Fig. 6C). We further subsampled the CD8^+^ T cell subset and thereby revealed that CD8^+^ T cells with a more naïve phenotype are enriched at the outer regions of the tumor, while exhausted CD8^+^ T cells mainly localize deeper inside the TME (Fig. 6D). Subsampling the monocyte/macrophage subset revealed a distinct localization pattern for different subsets of macrophages; namely, *Retnla^HI^* macrophages were exclusively found at the outer regions of the TME, while *Apoe^HI^*, *Ms4a7^HI^* and proliferating macrophages are skewed towards the interior of the TME (Fig. 6E).

In line with our previous data obtained using the *MMTV-PyMTChOVA* mouse model for spontaneous breast cancer (Hu et al., 2020), we found that CD8^+^ T cells show an increased exhaustion score (Wherry et al., 2007) when located in the inner regions of the TME (Fig. 6F, left). Moreover, we applied pseudotime analysis (Cao et al., 2019) on the monocyte/macrophage subset, and specified the *Ly6c2^HI^* monocyte-like cells as the root state of the trajectory which resulted in the *Apoe^HI^*, *Ms4a7^HI^* and proliferating macrophages as the terminally states (Fig. S6C, D), consistent with our previous findings (Hu et al., 2020; Mujal et al., 2021). When we overlaid Zipseq spatial localization on our pseudotime trajectory, we found a correlation with pseudotime progressing from monocyte-like early states at the outer regions to terminally differentiated TAM states in the tumor core (Fig S6E). This was also reflected in an advanced monocyte/macrophage pseudotime score when moving from outer towards the interior of the tumor (Fig. 6G, left). Interestingly, the positive correlation between the expression of exhaustion-related genes in CD8^+^ T cells and macrophage maturation towards the inner regions of the TME coincided with an increased glycolytic score (Argüello et al., 2020) in CD8^+^ T cells, but not in the monocyte/macrophage fraction (Fig. 6F, right and Fig. 6G, right), consistent with a more hypoxic microenvironment which correlates with this dysfunctional crosstalk between T_ex_ and TAM.

To predict interaction likelihood between different cell types in distinct regions in the TME, we used CellChat analysis, which uses a curated database of receptor-ligand interactions to highlight likely cell-cell interactions (Jin et al., 2021). These analyses revealed that expression of the CSF1-CSF1R ligand-receptor pair is significantly enriched in likelihood, and especially in the inner region versus outer region of tumors (Fig. 6H). Interestingly, CSF1 is found to be exclusively expressed by T_ex_ in those regions (Fig. 6H). CellChat analysis also predicted that the main receivers of T_ex_-derived CSF1 in the inner regions of the tumor are *Ms4a7^HI^* macrophages, proliferating macrophages and *Apoe^HI^* macrophages (Fig. 6H), pointing towards a co-dependency between TAM and T_ex_. In line with this, we found a positive correlation between an *‘exhaustion’* signature, as well as normalized *Csf1* and *Ccl4* expression in CD8^+^ T cells and macrophage maturation (monocyte/macrophage pseudotime score), when moving from the outer towards the inner regions of the tumor (Fig. 6I). Conversely, expression of genes associated with antigen presentation in monocyte/macrophages gradually decreased when moving closer towards the inner regions of the tumor (Fig. 6I). These data support a model in which monocytes, as they move inward and differentiate toward terminal TAM, downregulate antigen presentation in concert with the development of the exhausted state in T cells.

## Discussion

We mechanistically dissected a cellular co-alignment by which tumor-associated macrophages (TAM) and exhausted CD8^+^ T cells (T_ex_) in the TME co-exist in a self-enforcing positive feedback loop in mouse and human cancers. This includes finding that the secretion of growth factors and chemokines by one induces the other, the key interaction biology — a weakly stimulatory, yet long-duration synapse that ‘primes’ T cells for exhaustion — and spatial transcriptomics that demonstrate the co-evolution of these differentiated cell states, across space, in tumor tissue. Together, this demonstrates a principle of co-evolution of immunosuppressive cell types in the TME that supports immune evasion rather than destruction of the tumor.

The presence of TAM in solid tumors often correlates with poor prognosis and failure of response to anti-cancer therapies (Zhang et al., 2012; De Palma and Lewis, 2013). These findings are consistent with the established role of TAM in suppressing anti-tumor T cell immunity (DeNardo and Ruffell, 2019), and recent single cell RNA sequencing studies and other immune profiling approaches have hinted towards a potential link between the presence of TAM and exhausted CD8^+^ T cells in several different cancer types (Bi et al., 2021; Braun et al., 2021; Combes et al., 2021; Hong et al., 2021; Hu et al., 2020; Mujal et al., 2021; O’Connell et al., 2021; Wagner et al., 2019). However, these computational predictions still require experimental investigation to establish causality. Building on previous findings from our own lab and others that CD8^+^ T cells preferentially localize in TAM-rich areas in the TME (Boissonnas et al., 2013; Boldajipour et al., 2016; Broz et al., 2014; Engelhardt et al., 2012; Peranzoni et al., 2018), we show here that the evolution of these long considered immunosuppressive cell types in the TME is extensively linked in a causal circuit.

Macrophages are known to display a remarkable heterogeneity and plasticity that is dependent on a variety of environmental cues, some of which are derived from T cells (DeNardo and Ruffell, 2019; Guerriero, 2019). However, prior to our study, a role for T_ex_ in shaping macrophage phenotype and function has not been reported. We find in our models that intratumoral T_ex_ are the main immune population producing CSF1 to actively recruit monocytes and modulate their differentiation trajectory favoring antigen-presentation. Thinking beyond these models, other cell types in the TME, including tumor cells and fibroblasts, can also modulate myeloid biology through secretion of CSF1 (Buechler et al., 2021), and so there may be settings in which additional cell types also contribute to the establishment of the TAM-T_ex_ axis. However, in our data CSF1 is a predominant and consistent ‘exhaustion’ gene, and indeed previous reports have identified CCL3, 4 and 5 as amongst the most differentially expressed genes in chronic viral infection-associated exhaustion (Wherry et al., 2007). We consider it likely that T_ex_ express additional myeloid-modulating factors that have yet to be identified and may now be sought, based on these studies.

Computational analysis of our spatial ZipSeq data suggests that T_ex_-derived CSF1 is most likely to affect specific subpopulations of terminally differentiated macrophages that are enriched in the inner regions of the TME, including Ms4a7^HI^, Apoe^HI^, and proliferating TAM. While other scRNAseq studies have also reported on the existence of multiple different subpopulations of monocytes and TAM in the TME (Hu et al., 2020; Katzenelenbogen et al., 2020; Molgora et al., 2020; Mujal et al., 2021), it remains to be determined whether these are functionally distinct from one another in their interactions with CD8^+^ T cells and their abilities to modulate the onset of T cell exhaustion.

Regardless of their ability to phagocytose large amounts of antigen, TAM are often considered inferior in antigen processing and presentation as compared to conventional dendritic cells. Here we show that antigen-presenting TAM capture CD8^+^ T cells in uniquely long-lasting synaptic interactions characterized by the formation of variegated TCR microclusters. Despite expressing similar levels of MHC class I and II, co-stimulatory molecules and genes involved in cross-presentation as do cDC1 (Broz et al., 2014), TAM trigger only a weak TCR stimulation that fails to support proliferation, but clearly primes the onset of T cell exhaustion which is not observed in the absence of TCR-ligands presented by these TAM. Notably, blockade of immune checkpoint molecules PD1/PD-L1 and CTLA4 is unable to license proliferation in T cells responding to TAM (Engelhardt et al., 2012). It is clear that more work is required to better understand the fundamental nature of disparate TCR triggers and co-stimulation over time that contribute to the hyporesponsive state in T cells during tumorigenesis, as elegantly reviewed recently (Philip and Schietinger, 2021).

Recent studies have reported that development of T cell exhaustion during chronic infection and cancer occurs in a multistep fashion, revealing distinct subtypes with unique transcriptional and epigenetic dynamics, as well as their ability to respond to immune checkpoint blockade (Im et al., 2016; Philip et al., 2017; Satpathy et al., 2019; Siddiqui et al., 2019; Jansen et al., 2019; Miller et al., 2019; Beltra et al., 2020; Pritykin et al., 2021). The decision-making during this bifurcative process seems to be tightly regulated by transcription factors like IRF4 (Utzschneider et al., 2020; Chen et al., 2021; Seo et al., 2021) which we find is strongly upregulated by TAM despite only subtle other signs of TCR engagement. Furthermore, our *in vitro* co-culture studies suggest that TAM at least are capable of mediating the early stages of exhaustion, which is exacerbated in hypoxic conditions. Hypoxia was recently shown to be an important co-factor in the induction of exhaustion *in vitro* (Scharping et al., 2021), and ZipSeq transcriptomics here places terminal exhaustion *in vivo* preferentially in hypoxic regions of the TME. In line with this, other recent studies have elegantly demonstrated how metabolic insufficiencies drive mitochondrial stress in T cells contributing to exhaustion phenotypes in chronic infections and cancer (Bengsch et al., 2016; Thommen et al., 2018; Vardhana et al., 2020; Scharping et al., 2021).

Taken together, our work dissects a novel spatiotemporal co-evolution between exhausted CD8^+^ T cells and TAM in the TME that supports tumor evasion rather than tumor destruction. We believe that co-dependency of different lineages may explain some of the resistance of the TME to targeting — removing just one cell population will still leave the other, to influence re-establishment of the targeted one. Thus, therapeutic strategies may need to break the biology underlying the TAM-T_ex_ axis at multiple points. Doing so may work in conjunction with existing immunotherapies to enhance anti-tumor immunity, and thereby expand the proportion of cancer patients who benefit from immunotherapies.

## Acknowledgements

We thank all members of the Krummel laboratory for discussion and support. We also thank the ImmunoX CoLabs at UCSF for technical assistance and support. Figures were created using BioRender.com. Funding: K.K. was supported by a Rubicon postdoctoral fellowship 019.163LW.006 from the Netherlands Organization for Scientific Research (NWO), and a Parker Scholar Award from the Parker Institute for Cancer Immunotherapy. K.H.H. was supported by a Parker Scholar Award from the Parker Institute for Cancer Immunotherapy, and a Jean Perkins postdoctoral fellowship from the American Cancer Society. A.R. and E.C. were supported by postdoctoral fellowships from the Cancer Research Institute. J.A.B was supported by a Stanford Graduate Fellowship and the National Science Foundation Graduate Research Fellowship under Grant No. DGE-1656518. A.T.S. was supported by NIH U01CA260852, a Cancer Research Institute Technology Impact Award, and a Career Award for Medical Scientists from the Burroughs Wellcome Fund. This work was supported by NIH/NCI U54CA163123, 5U01CA217864, R21CA191428 and R01CA197363 to M.F.K.

## Author contributions

K.K. designed and conducted most experiments, data analysis, and drafted the manuscript. K.H.H. and A.R. designed, performed and analyzed ZipSeq experiments. A.J.C. and B.S. analyzed human cancer patient datasets. T.H. assisted with co-culture experiments. A.A.R. analyzed RNAseq data. E.C., K.M. and J.A. provided technical support and assisted with imaging experiments. T.C. discussed data and project direction. Q.S., J.A.B. and A.T.S. performed ATACseq and assisted with data analysis. M.F.K. designed the experiments, interpreted data, and with other co-authors, developed the manuscript.

## STAR Methods

### Key Resource Table

See Excel Key Resource Table.

### Contact for Reagent and Resource Sharing

Further information and requests for resources and reagents should be directed to and will be fulfilled by the Lead Contact, Matthew F. Krummel.

### Experimental Model and Subject Details

#### Human tumor samples

Flow cytometry on RCC samples: RCC were transported from various cancer operating rooms or outpatient clinics. All patients consented by the UCSF IPI clinical coordinator group for tissue collection under a UCSF IRB approved protocol (UCSF IRB# 20-31740). Samples were obtained after surgical excision with biopsies taken by Pathology Assistants to confirm the presence of tumor cells. Patients were selected without regard to prior treatment. Freshly resected samples were placed in ice-cold DPBS or Leibovitz’s L-15 medium in a 50 mL conical tube and immediately transported to the laboratory for sample labeling and processing. The whole tissue underwent digestion and processing to generate a single-cell suspension. In the event that part of the tissue was sliced and preserved for imaging analysis, the remaining portion of the tissue sample was used for flow cytometry analysis as described in Combes et al (Combes et al., 2021).

Samples from the following indications were used for RNAseq on FACS-isolated cell fractions performed as described previously (Combes et al., 2021): Bladder cancer (BLAD), colorectal cancer (CRC), glioblastoma multiforme (GBM), endometrial and ovarian cancer (GYN), hepatocellular carcinoma (HEP), head and neck squamous cell carcinoma (HNSC), (KID), lung adenocarcinoma (LUNG), skin cutaneous melanoma (MEL), pancreatic ductal adenocarcinoma (PDAC), pancreatic neuroendocrine tumors (PNET), sarcoma (SRC).

#### Mice

All mice were treated in accordance with the regulatory standards of the National Institutes of Health and American Association of Laboratory Animal Care and were approved by the UCSF Institution of Animal Care and Use Committee. The following mice were purchased for acute use or maintained under specific pathogen-free conditions at the University of California, San Francisco Animal Barrier Facility: C57BL6/J, C57BL6/J CD45.1, OT-I, P14 LCMV, Rag1-/-, CSF1op/op, mTmG. With the exception of CSF1op/op, all mice used in experimentation were bred to a C57BL6/J background. Mice of either sex ranging in age from 6-12 weeks were used for experimentation. For experiments using the transgenic MMTV-PyMTChOVA strain (Engelhardt et al., 2012), only mammary tumor-bearing females were used ranging in age from 12-20 weeks. Treatments in MMTV-PyMTChOVA mice were started when mammary tumors reached ∼25mm2 in size. CSF1RCreCD206-LSL-Venus-DTR mice (Ray, A. et al. in preparation) were generated and used for ZipSeq. Food and water were provided ad libitum.

#### Tumor cell lines

Tumor cell lines B16F10 (CRL-6475, ATCC), B16ChOVA (Roberts et al., 2016; Binnewies et al., 2019) and B78ChOVA (Engelhardt et al., 2012; Broz et al., 2014) were cultured under standard conditions 37°C in 5% CO2 in DMEM (GIBCO), 10% FCS (Benchmark), 1% Pen/Strep/Glut (Invitrogen).

### Method Details

#### Tumor growth experiments

For tumor studies, adherent tumor cells were grown to confluency and harvested using 0.05% Trypsin-EDTA (GIBCO) and washed 3x with PBS (GIBCO). 1.0×105 – 2.5×105 cells in PBS were resuspended in a 1:1 ratio with Growth Factor Reduced Matrigel (Corning) and a final volume of 50µl was injected subcutaneously into the flanks of anaesthetized and shaved mice. Tumors were allowed to grow for 14-21 days unless otherwise noted, before tumors and tumor-draining lymph nodes were harvested for analysis.

#### Adoptive T cell transfers

Inguinal, axillary, brachial and mesenteric lymph nodes (LN) or spleens were isolated from CD45.1 OT-I or P14 LCMV mice. LN and spleens were meshed through 70µm filters and treated with ACK red blood cell lysis buffer. CD8+ T cells were purified using EasySep CD8 negative selection kits (Stemcell Technologies). 1×105 (for >14 day read-out) – 2×106 T cells (for day 4 read-out) were adoptively transferred through retro-orbital injection in 100µl PBS.

For the comparison of OT-I T cells and p14 LCMV T cells, mice received a 1:1 mix of both T cells in 100µl of PBS through retro-orbital injection. The following day mice were inoculated with a bolus of CFA containing gp33-peptide (50µg/mouse; Anaspec) and SL8/SIINFEKL peptide (50µg/mouse; Anaspec) subcutaneously, to sustain both T cell populations.

#### In vivo antibody treatment

For macrophage depletions, mice received anti-CSF1 (clone 5A1; BioXCell), anti-CSF1R (clone AFS98; BioXCell) or corresponding isotype controls, Rat IgG1k (clone HRPN; BioXCell) and Rat IgG2a (clone 2A3; BioXCell), respectively. Antibodies were injected intraperitoneally at an initial dose of 1mg/mouse followed by 0.5mg/mouse every 7 days.

For T cell depletion studies, mice received anti-CD4 (clone GK1.5; BioXCell), anti-CD8a (clone 2.43; BioXCell) or corresponding isotype control, Rat IgG2b (clone LTF-2; BioXCell) dosed at 250 µg/mouse every 3-4 days.

#### Generation of mixed bone marrow chimeras

Mixed bone marrow chimeras were generated as described previously (Barry et al., 2018). Briefly, Rag1-/- mice were lethally irradiated with 1,100 rads of irradiation in two doses 3-5 hours apart. 2-5×106 bone marrow cells, consisting of 50% Rag1-/- and 50% CSF1op/op or CSF1op/+ bone marrow, were injected retro-orbitally to reconstitute irradiated mice. Chimeric mice were allowed to recover for 6-10 weeks, upon which mice were inoculated with B78ChOVA tumors subcutaneously.

#### Mouse tissue digestion and flow cytometry

Tumors were harvested and processed to single cell suspensions as described previously (Barry et al., 2018; Binnewies et al., 2019). Briefly, tumors were isolated and mechanically minced, followed by enzymatic digestion with 200µg/ml DNAse (Sigma-Aldrich), 100U/ml Collagenase I (Worthington Biochemical) and 500U/ml Collagenase Type IV (Worthington Biochemical) for 30 minutes at 37°C while shaking. Enzymatic activity was quenched by adding equal amounts of FACS buffer (2% FCS in PBS), and cell suspensions were filtered to obtain single cell suspensions. TdLN were isolated and meshed over 70µm filters in PBS to generate single cell suspensions. For each sample, 5-10×106 cells were used for staining for flow cytometry. Cells were washed with PBS prior to staining with Zombie NIR Fixable live/dead dye (Biolegend) for 20 min at 4°C. Cells were washed in PBS followed by surface staining for 30 min at 4°C with directly conjugated antibodies diluted in FACS buffer containing anti-CD16/32 (BioXCell) to block non-specific binding. Cells were washed again with FACS buffer. For intracellular staining, cells were fixed for 20 min at 4°C using the FOXP3 Fix/Perm kit (BD Biosciences), and washed in permeabilization buffer. Antibodies against intracellular targets were diluted in permeabilization buffer and cells were incubated for 30 min at 4°C followed by another wash prior to read-out on a BD LSR Fortessa SORP cytometer.

#### Fluorescence Activated Cell Sorting

Single cell suspensions from tumors were prepared as described above. For T cell isolations, single cell suspensions were enriched for mononuclear cells using Ficoll-Paque Premium 1.084 (GE Healthcare). For isolation of myeloid cells, single cell tumor suspensions were enriched for CD45+ cells using EasySep biotin positive selection kit (Stemcell Technologies). Enriched cells were stained for 30 min at 4°C with directly conjugated antibodies diluted in FACS buffer containing anti-CD16/32 (BioXCell) to block non-specific binding. Cells were washed again with FACS buffer and filtered over a 70µm mesh. Immediately prior to sorting, DAPI was added to exclude dead cells. Cells were sorted on a BD FACSAria Fusion and BD FACSAria2. Sorted T cells were collected directly in lysis buffer (Invitrogen) for RNA sequencing or in RPMI (GIBCO), 10% FCS (Benchmark), Pen/Strep/Glut (Invitrogen) and 50µM *β*-mercaptoethanol (GIBCO) at 4°C for further use ex vivo. Sorted myeloid cells were collected in DMEM (GIBCO), 10% FCS (Benchmark), Pen/Strep/Glut (Invitrogen) at 4°C for further use ex vivo.

#### T cell cytokine analysis

For analysis of cytokine production by endogenous and adoptively transferred T cells, 5-10×106 LN and tumor cells were re-stimulated for 3-5 hours in RPMI (GIBCO), 10% FCS (Benchmark), Pen/Strep/Glut (Invitrogen), 50µM *β*-mercaptoethanol (GIBCO) containing PMA (50ng/ml; Sigma-Aldrich), ionomycin (500ng/ml; Invitrogen) and brefeldin A (3µg/ml; Sigma-Aldrich) at 37°C in 5% CO2. Cells were washed and stained for intracellular flow cytometric analysis.

#### RNA sequencing

mRNA from cells were isolated using DynaBead Direct and then converted into amplified cDNA using the Tecan Ovation RNA-Seq System V2 kit, following the manufacturer guidelines. The dsDNA is tagmented, amplified and undergoes clean up with AMPure XP bead, using the Illumina Nextera XT DNA Library Prep Kit. The resulting sequencing library is QC’d using an Agilent Bioanalyzer HS DNA chip to assess fragment size distribution and concentration. Libraries were pooled prior to single-end sequencing on and Illumina MiSeq/MiniSeq to ensure quantify library complexity. Libraries with less than 10 percent of the reads aligned to coding regions, or fewer than 1,000 unique reads in total were rejected. The validated libraries were re-pooled based on the percentage of reads in coding regions and submitted to the UCSF Center for Advanced Technology for 150bp paired end sequencing on an Illumina NovaSeq 6000.

Raw fastq reads were QC’d and trimmed to remove adapter contamination, and poly-G artifacts using using fastp version 0.19.6 (Chen et al., 2018). Reads with fewer than 20bp post-trimming were discarded. Trimmed reads were aligned to the GRCm38 reference sequence annotated with Gencode V25 (Frankish et al., 2019) using STAR version 2.6.1b (Dobin et al., 2013) with the following parameters (--quantMode GeneCounts–outFilterMismatchNoverLmax 0.04 --alignIntronMax 100000 --alignMatesGapMax 100000 -- alignSJDBoverhangMin 10 --alignSJstitchMismatchNmax 5 -1 5 5 --chimSegmentMin 12 -- chimJunctionOverhangMin 12 --chimSegmentReadGapMax 3 --chimMultimapScoreRange 10 -- chimMultimapNmax 10 --chimNonchimScoreDropMin 10 --peOverlapNbasesMin 12 --peOverlapMMp 0.1) STAR-generated reads counts from each library were processed using the limma/Voom pipeline (Law et al., 2014; Smyth, 2005) using the edgeR package (Robinson et al., 2010). Briefly, the read counts are loaded into a DGEList object to generate Counts Per Million (CPM), and then filtered to retain only genes with at least 10 counts in a worthwhile number of samples and at least 15 counts across all samples. The CPM matrix is normalized using TMM Trimmed mean of M-values and processed using voom to estimate the mean-variance relationship to identify edge weights that can be used to fit to a linear model with limma lmFit. Differential gene expression between two groups of empirical Bayes moderation of the standard errors towards a global value. A list of transcriptional DEGs between Tnaïve and Tex d14 with a FC equal to or >5 was generated and gene set enrichment analysis was performed using the MGI Mammalian Phenotype Level 4 database in Enrichr (Chen et al., 2013; Kuleshov et al., 2016; Xie et al., 2021).

#### ATAC sequencing

ATAC-seq samples were processed according to the Omni-ATAC protocol (Corces et al., 2017). 5×104 cells per replicate were lysed in 50 µL ATAC resuspension buffer supplemented with 0.1% NP40, 0.1% Tween-20, and 0.01% Digitonin. After lysis, nuclei were transposed using 2.5 µl Tn5 transposase in a 50 µl reaction for 30 min at 37°C. Finally, the transposed DNA was purified using a commercial PCR cleanup kit and libraries were prepared for sequencing. 2×75 paired end sequencing was performed on an Illumina sequencer.

ATAC-seq computational analysis was performed as previously described (Weber et al., 2021). Briefly, read trimming and filtering was performed with fastp. Reads were mapped to the hg38 reference genome using hisat2 with the --no-spliced-alignment option. Picard was used to remove duplicates from bam files. We removed any reads not mapping to chromosomes 1-22 and chrX (ie chrY reads, mitochondrial reads, and other reads were discarded). The deduplicated and filtered fragments were then formatted into a bed file. Peaks were called using MACS2. Peaks from each sample were iteratively merged into a high confidence union peak set for all samples as previously described (Corces et al., 2018). A peak by sample matrix was created by overlapping fragments in each sample with each peak, and this matrix was used to perform differential peak analysis in DESeq2. Genome coverage files were created from the fragments file by loading the fragments into R and then exporting bigwig files normalized by reads in transcription start sites using ‘rtracklayer::export’. Normalized track files were visualized using the Integrative Genomics Viewer.

#### qRT-PCR

RNA was extracted from FACS-sorted immune cell populations using Qiagen RNeasy Micro kit (Qiagen) and the yield was measured using Nanodrop. cDNA first-strand synthesis was performed using High-Capacity cDNA Reverse Transcription kit (Thermo Fisher Scientific) using random primers. qRT-PCR analysis was performed using Taqman probes targeting Ccl3 (Mm00441259_g1), Ccl5 (Mm01302427_m1), Csf1 (Mm00432686_m1) and Gapdh (Mm99999915_g1). All probes were obtained from Life Technologies. For amplification reactions, iTaq Universal Probes Supermix was used according to manufacturer’s instructions. qRT-PCR was performed on a QuantStudio 12K Flex lightcycler (Applied Biosystems by Life Technologies). For quantification the delta Ct method was used: delta Ct sample — delta Ct reference gene. All transcripts were normalized to Gapdh.

#### Monocyte recruitment transwell assays

Bone marrow was obtained from femurs and tibia of CD45.1 mice, and monocytes were isolated using EasySep Mouse Monocyte Isolation kits (Stemcell Technologies). For transwell assays, 1×105 monocytes were added to top inserts containing 5.0 µm pore polycarbonate membrane (Corning). 0.5×105 naïve, previously activated or exhausted OT-I T cells were cultured in bottom wells in RPMI (GIBCO), 10% FCS (Benchmark), Pen/Strep/Glut (Invitrogen) and 50µM *β*-mercaptoethanol (GIBCO). Migration through the membrane was analyzed after 24 hours of culture. Plate was briefly centrifuged briefly at 1000rpm for 1 min to collect cells stuck to the membrane. Cells were collected for analysis by flow cytometry. Absolute counting beads (Life Technologies) were added for quantification of the number of migrated cells.

#### Generation of activated T cells

OT-I T cells were activated in vitro as described previously (Broz et al., 2014). Briefly, OT-I lymph node cells were stimulated with B6 splenocytes pulsed with SL8 peptide (100ng/ml; Anaspec) for 30 min at 37°C and then washed 3 times. On day 2-3, cells were expanded by adding human IL-2 (2U/ml; Peprotech) to fresh RPMI (GIBCO), 10% FCS (Benchmark), Pen/Strep/Glut (Invitrogen), 50µM *β*-mercaptoethanol (GIBCO). Cells were used for co-culture assays on day 4-5. Prior to use dead cells were excluded using Ficoll-Paque PLUS (GE Healthcare).

#### Bone marrow-derived dendritic cells

BMDC were generated as described previously (Broz et al., 2014). Briefly, bone marrow was obtained from femurs and tibia of C57BL6/J mice and cultured in DMEM (GIBCO), 10% FCS (Benchmark), Pen/Strep/Glut (Invitrogen) in the presence of 7.5 ng/ml GM-CSF (Peprotech) for 6-8 days, followed by the addition of 60ng/ml IL4 (Peprotech) for the last 2 days. Media was refreshed every 3-4 days. For co-cultures studies, BMDC were pulsed with SL8 peptide (100ng/ml; Anaspec) for 30 min at 37°C and then washed 3 times prior to use.

#### Quantification of APC-T cell interactions

APC (BMDC, BMDC+SL8 or sorted TAM) were obtained from mTmG mice, and previously activated OT-I T cells were stained for 15 minutes at 37°C with 2 µM CFSE (Invitrogen) in PBS and washed in RMPI prior to use. Cells were co-cultured in NUNC 8 well chamber slides (Thermo Scientific) that were coated with fibronectin (2µg/ml; EMD Millipore) in PBS at 37°C for 1 hour before use. APCs in phenol red-free RPMI were allowed to attach to the chamber slides for 20-30 minutes at 37°C and 5% CO2. Right before imaging, T cells (resuspended in 0.1% agarose) were added to the wells and slides were loaded for imaging. To visualize the interaction between different APC populations and T cells, a conventional widefield Zeiss Axiovert 200M was used with a Sutter Lambda XL illumination source, running on µMagellan software. Images were acquired every 2 minutes for 6 hours using a 20x objective. Samples were kept at 37°C using a heated robotic stage. Image analysis was performed in Imaris (Bitplane) and ImageJ.

#### Lattice light-sheet imaging

Lattice light-sheet (LLS) imaging was performed in a manner previously described (Cai et al., 2017). Briefly, 5 mm diameter round coverslips were cleaned by a plasma cleaner and coated with fibronectin (2µg/ml; EMD Millipore) in PBS at 37°C for 1 hour before use. TAM sorted from B78ChOVA-bearing mTmG mice were dropped onto the coverslip and incubated at 37°C, 5% CO2 for 20–30 min. Previously activated OT-I CD8+ T cells were labeled with CD45-AF647 (clone 30-F11; Biolegend) and TCR*β* AF488 (clone H57-597; BioLegend) for 30 min and washed in FACS buffer. Right before imaging, T cells were dropped onto the coverslip containing TAM. The sample was then loaded into the previously conditioned sample bath and secured. Imaging was performed with a 488-nm, 560-nm, or 642-nm laser (MPBC, Canada) dependent upon sample labeling in single or two-color mode. Exposure time was 10 ms per frame leading to a temporal resolution of 4.5 s. Image renderings were created using Imaris software (Bitplane). Quantification of TCR clustering was performed using ImageJ. Briefly, channels were separated and entire T cell membrane versus TAM interaction site was outlined manually. Signal intensity for red (CD45) and green (TCR) channel was calculated. The following formula was used to determine TCR signal intensity for both channels at the synaptic TAM-T cell interaction site: signaling intensity = (intensity at synapse/intensity total membrane).

#### FURA-2AM Calcium imaging

TAM or CD103+ DC were sorted from B16ChOVA or B16F10 tumors as per description above, and stained with 2 µM CMTMR (Thermo Fisher Scientific) in PBS for 15 minutes at 37°C, followed by a wash in RPMI. Cells were co-cultured in NUNC 8 well chamber slides (Thermo Scientific) that were coated with fibronectin (2µg/ml; EMD Millipore) in PBS at 37°C for 1 hour before use. APCs in phenol red-free RPMI were allowed to attach to the chamber slides for 20-30 minutes at 37°C and 5% CO2. Right before imaging, previously activated OT-I T cells were labeled with FURA-2 AM (0.5µM; Invitrogen) for 15 minutes at RT. Cells were washed and resuspended in phenol red-free RPMI (GIBCO), 10% FCS (Benchmark), Pen/Strep/Glut (Invitrogen), 50µM *β*-mercaptoethanol (GIBCO) supplemented with 0.1% agarose and were added to the wells. Imaging was performed using an Zeiss Axiovert 200M microscope with a Sutter Lambda XL illumination source equipped with a 40x oil objective. Images were acquired every 5 seconds for 18-21 minutes. Samples were kept at 37°C using a heated robotic stage. Image analysis was performed in Imaris (Bitplane). Briefly, background subtraction was performed and surfaces were created for APCs and T cells to quantify dwell time to determine whether cells were touching (cut-off equal to or >3). Calcium2+ flux was determined by calculating the average 340/380 ratiometric fluorescence per cell after contact for each time point.

#### Hypoxyprobe imaging

Mice were injected with pimonidazole hydrochloride in PBS (80mg/kg; Hypoxyprobe) intraperitoneally 1.5 hours prior to sacrifice. Tissues were dissected and processed as described above. For flow cytometry studies, pimonidazole was visualized using anti-pimonidazole antibodies (Pacific Blue Mab-1 clone 4.3.11.3; Hypoxyprobe) after cells were fixed for 20 min at 4°C using the FOXP3 Fix/Perm kit (BD Biosciences), and washed in permeabilization buffer. For imaging studies, dissected tumors were embedded in OCT and sectioned into 10µm cryosections. Cryosections were stored at -80°C until further use. For immunostaining, sections were fixed in 4% PFA (Electron Microscopy Sciences) for 20 minutes at RT, followed by a rinse in PBS containing 1% BSA (Sigma). Sections were blocked in 1% BSA in PBS containing anti-CD16/32 (BioXCell) for 1 hour at RT, and washed. Sections were stained with CD11b-AF594 (clone M1/70;BioLegend), CD31-AF647 (clone 390;BioLegend) and Pacific Blue Mab-1 (clone 4.3.11.3;Hypoxyprobe) for 1 hour at RT, followed by a wash and mounted using Vectashield (Vector Laboratories) and sealed with nail polish. Images were acquired on a Leica SP8 confocal microscope. Data analysis was performed using Imaris (Bitplane).

#### In vitro APC-T cell co-culture assays

APC populations were sorted from tumors as described above, and co-cultured with 1×105 previously activated OT-I CD8+ T cells labeled with Violet Proliferation dye (VPD; BD Biosciences) at a 1:5 ratio (unless otherwise noted) in RPMI (GIBCO), 10% FCS (Benchmark), Pen/Strep/Glut (Invitrogen), 50µM *β*-mercaptoethanol (GIBCO) in 96-well round bottom plates. Cells were harvested for analysis 3 days later, unless otherwise noted. In vitro generated BMDC and BMDC+SL8 were used as negative and positive controls, respectively.

For hypoxia experiments, plates containing exact same experimental groups were incubated in a Avatar hypoxic bioreactor (XcellBio) at 1.5% O2 or under ambient 21% O2 for comparison and cells were harvested after 3 days.

#### ZipSeq

ZipSeq spatial transcriptomics was performed as described previously (Hu et al., 2020). Briefly, B78ChOVA cells were injected subcutaneously as described above and were harvested day 16 post-injection. Tumors were sectioned while live using a compresstome (Precisionary Instruments VFZ-310-0Z) to generate ∼160 µm sections. The sectioning, imaging, spatial barcoding, tumor dissociation, sorting, 10X encapsulation and library construction were identical to the methods described in (Hu et al., 2020). The targeted number of cells for loading was 5000. With this in mind, we aimed for 30,000 reads per cell during sequencing on an Illumina S4 flowcell with a 1:10 molar ratio of Zipcode reads to gene expression reads. Resulting fastq files were processed using the CellRanger 4.0.0 pipeline, aligning to the GRCm38 Mus musculus assembly. CellRanger output thus resulted in ∼359k reads for the gene expression library and ∼40k reads for the Zipcode library.

#### Analysis of scRNA-Seq

The raw feature-barcode matrix generated by 10X CellRanger was loaded into Seurat (Satija et al., 2015). Cells with mitochondrial read % over 20% and those with less than 500 genes detected were excluded from analysis. Zipcode read counts from CellRanger were also loaded into Seurat as a separate ‘ADT’ assay and using CLR normalized counts, cells with either too few Zipcode reads or mixed Zipcode reads were also excluded from analysis. Following built-in Seurat methods for gene expression normalization and variance stabilization (Single Cell Transform (Hafemeister and Satija, 2019)), cells underwent one more round of clean-up, removing a small cluster of contaminating CD45– cells and another small cluster dominated by mitochondrial and ribosomal genes. This yielded 2765 cells. At this stage, the mean # of UMI’s and # of detected genes was: (25,566 and 4,199 respectively) while for the antibody derived Zipcode tags the mean UMI was 1,891 reads. At this stage, we also determined cluster identities using Seurat’s FindAllMarkers function performed on the log-normalized read counts which by default uses the Wilcoxon Rank Sum test.

For CellChat analysis, this cleaned object was fed into the CellChat workflow, using the built-in mouse ligand-receptor database, and a tri-mean thresholding for significance of interaction. The Seurat object was split into 3 sub-objects based on regional assignment and these 3 objects were separately analyzed using the CellChat workflow for multiple datasets and then merged. For signature score generation, we used Seurat’s built-in AddModuleScore function with gene lists for Glycolysis (Argüello et al., 2020), T cell exhaustion (Wherry et al., 2007), and Antigen Presentation (GO term 0048002) using 50 control features. Full gene lists can be found in the Extended Data. For pseudotime analysis, the monocyte/macrophage sub-object was passed into Monocle v3 without any changes to the UMAP dimensional reduction.

## Quantification and Statistical Analysis

### Statistical analysis

Unless specifically noted, all data are representative of *≥* 2 separate experiments. Experimental group assignment was determined by random designation. Statistical analyses were performed using GraphPad Prism software. Error bars represent ± standard error of the mean (S.E.M.) calculated using Prism. Specific statistical tests used were paired t-tests, unpaired t-tests, and one-way ANOVA unless otherwise noted. P-values <0.05 were considered statistically significant. Investigators were not blinded to group assignment during experimental procedures or analysis.

## Data and Software Availability

## Supplementary Figure Legends

**Figure S1.**
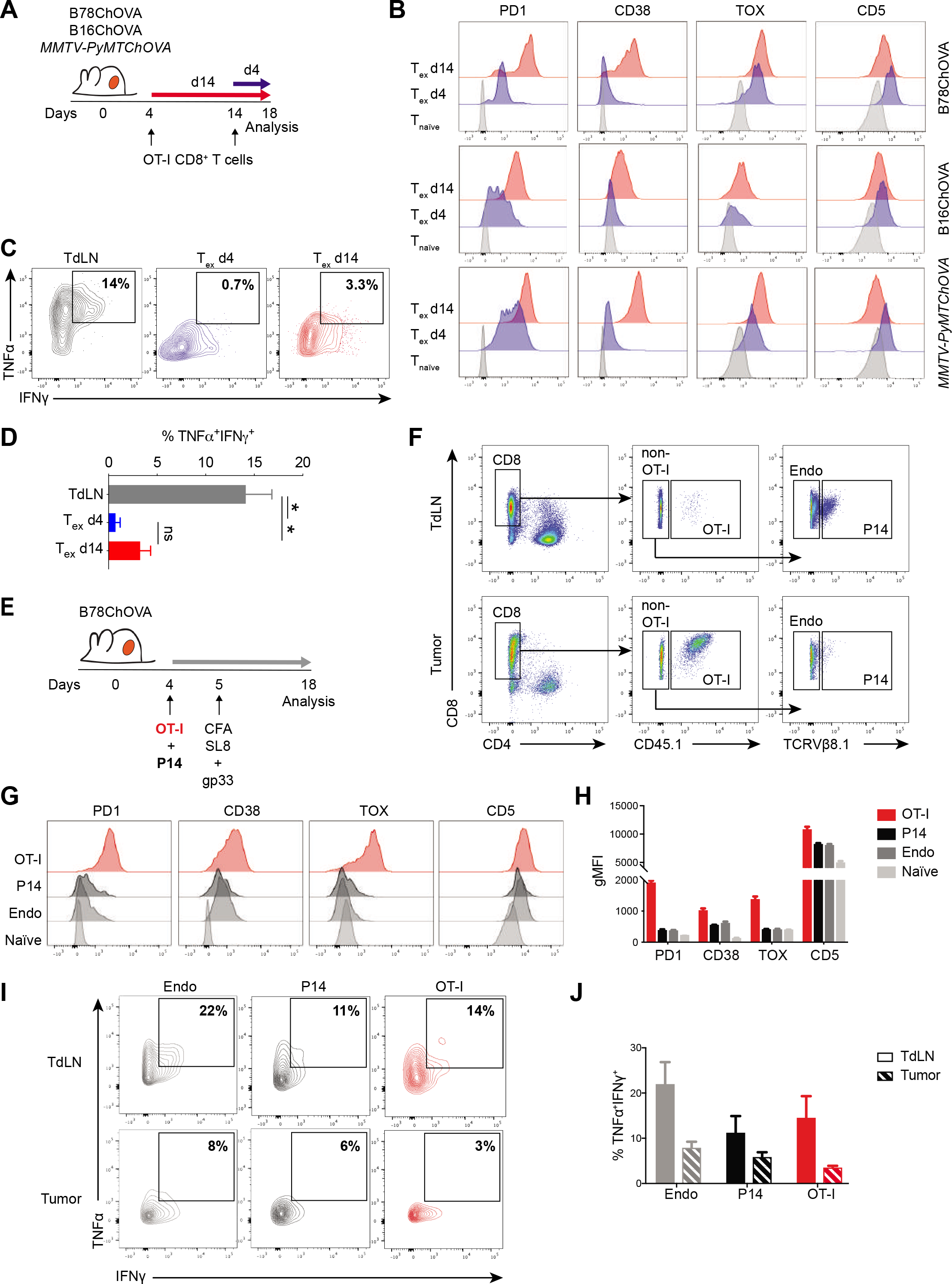
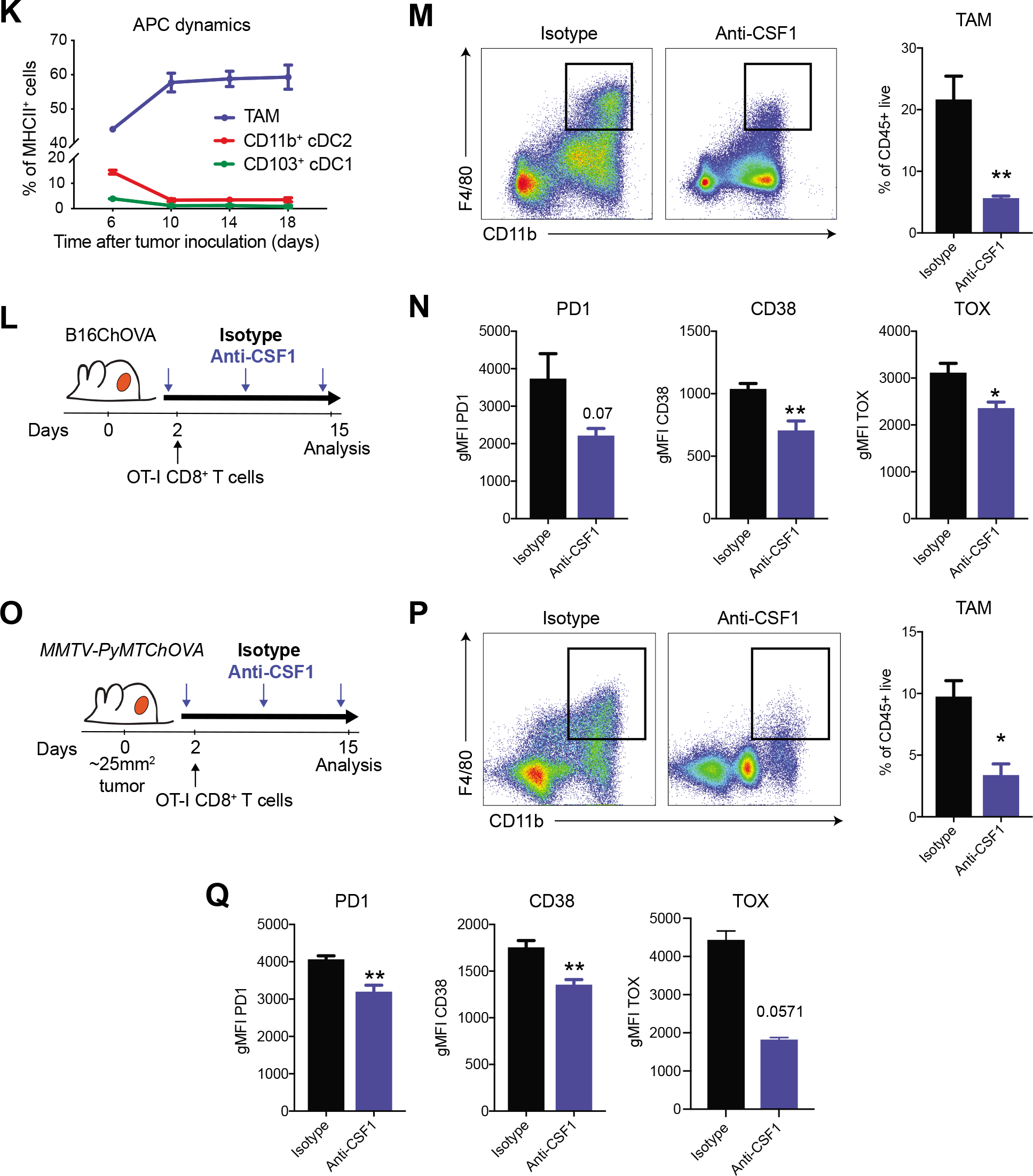
Onset of CD8^+^ T cell exhaustion is antigen-specific and correlates with macrophage abundance in multiple mouse cancer models (related to Figure 1). A) Experimental setup to study kinetics of CD8^+^ T cell exhaustion in B78ChOVA and B16ChOVA melanoma and spontaneous *MMTV-PyMTChOVA* breast cancer model. OVA-specific OT-I CD8^+^ T cells are adoptively transferred into tumor-bearing mice 14 days (T_ex_ d14) and 4 days (T_ex_ d4) prior to sacrifice, upon which tumors are harvested for analysis of T cell phenotype at day 18. B) Representative histograms of expression of PD1, CD38, TOX and CD5 expression on intratumoral CD44^+^ OT-I CD8^+^ T cells that have resided in the TME for 4 days (T_ex_ d4; blue) or 14 days (T_ex_ d14; red), versus naïve endogenous CD44^−^CD8^+^ T cells in the tumor-draining lymph node (TdLN) (T_naïve_). C-D) Representative contour plots (C) and quantification (D) of IFN*γ*^+^TNF*α*^+^ polyfunctional CD44^+^ OT-I CD8^+^ T cells that have resided for 4 days (T_ex_ d4) or 14 days (T_ex_ d14) in the TME compared to CD44^+^ endogenous CD8^+^ T cells in the TdLN (TdLN). N=3-10 mice/group. E) Experimental setup. Mice inoculated subcutaneously with B78ChOVA melanoma cells on day 0, received adoptively transferred OT-I and p14 LCMV CD8^+^ T cells i.v. on day 4, followed by inoculation with CFA containing SL8 + gp33 peptide s.c. on day 5. Mice were sacrificed on day 18 after tumor inoculation, and tumor-draining lymph nodes (TdLN) and tumors were harvested for analysis. F) Representative dot plots for the identification of endogenous (endo), and adoptively transferred CD45.1^+^ OT-I and TCRV*β*8.1^+^ P14 LCMV CD8^+^ T cells in TdLN (top) and tumors (bottom) by flow cytometry. G-H) Representative histograms (G) and quantification (H) of expression of PD1, CD38, TOX and CD5 on naïve CD44^−^CD8^+^ T cells in the TdLN and on tumor-infiltrating CD44^+^ endogenous (endo), P14 and OT-I CD8^+^ T cells. N = 5 mice/group. I-J) Representative contour plots (I) and quantification (J) of IFN*γ*^+^TNF*α*^+^ polyfunctional CD44^+^ endogenous (endo), P14 and OT-I CD8^+^ T cells in TdLN and tumor. N = 5 mice/group. Representative of two independent experiments. K) Quantification of TAM, CD11b^+^ cDC2 and CD103^+^ cDC1 populations represented as a fraction of MHCII^+^ cells in B78ChOVA tumors during tumor progression by flow cytometry. N=3 mice/time point. L) Experimental set-up of TAM depletion in B16ChOVA-bearing mice. Weekly anti-CSF1 treatment was initiated one day prior to adoptive transfer of OT-I CD8^+^ T cells. M) Representative dot plots and quantification of CD11b^+^ F4/80^+^ macrophages in isotype and anti-CSF1-treated B16ChOVA melanomas. N=5 mice/group. N) Surface (PD1 and CD38) and intracellular (TOX) expression on intratumoral CD44^+^ OT-I CD8^+^ T cells from isotype and anti-CSF1 treated B16ChOVA-bearing mice. N=5 mice/group. O) Experimental set-up of TAM depletion in spontaneous *MMTV-PyMTChOVA* breast cancer model. Weekly anti-CSF1 treatment was initiated when tumors reached ∼25mm^2^ in size and one day prior to adoptive transfer of OT-I CD8^+^ T cells. P) Representative dot plots and quantification of CD11b^+^ F4/80^+^ macrophages in isotype and anti-CSF1 treated mammary tumor-bearing *MMTV-PyMTChOVA* mice. N=5-6 tumors/group. Q) Surface (PD1 and CD38) and intracellular (TOX) expression on intratumoral CD44^+^ OT-I CD8^+^ T cells from isotype and anti-CSF1 treated mammary tumor-bearing *MMTV- PyMTChOVA* mice. N=3-6 tumors/group. All data are mean ± SEM. Statistical significance was determined using two-way ANOVA with Holm-Sidak’s correction for multiple comparisons or Mann-Whitney U test. * p < 0.05, ** p < 0.01, *** p < 0.001.

**Figure S2.**
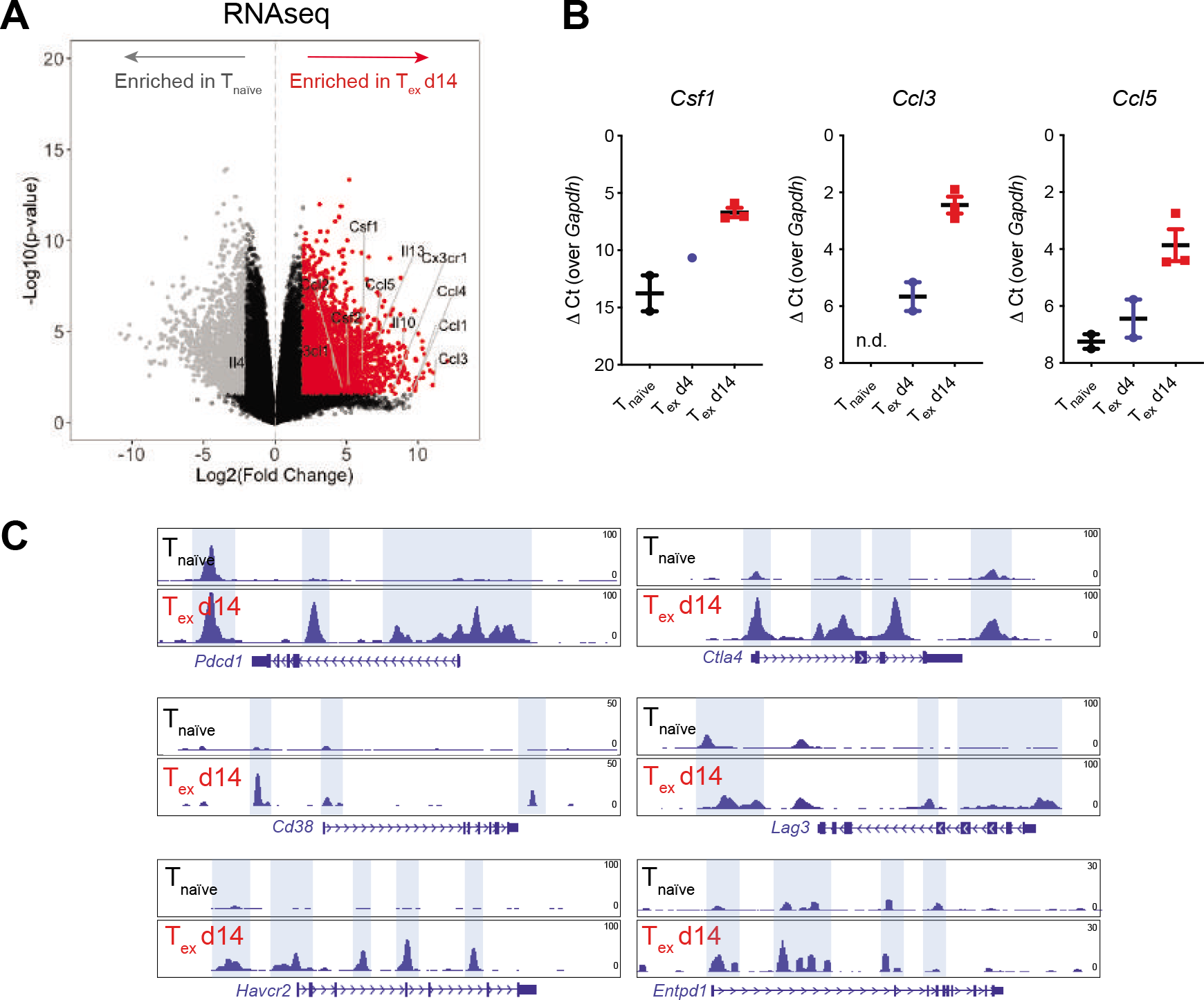
Transcriptional and epigenetic profiling reveals expression of myeloid-associated factors by CD8^+^ T_ex_ (related to Figure 2). A) Volcano plot showing differential gene expression in tumor-infiltrating CD44^+^ OT-I CD8^+^ T_ex_ d14 cells (red) compared to splenic CD44^−^ OT-I CD8^+^ T_naïve_ cells (grey) by RNAseq. Colored dots (grey and red) represent genes with a log2FC>2 and FDR<0.05. B) Expression of *Csf1*, *Ccl3* and *Ccl5* transcripts in an independent sample set of T_naïve_, OT-I T_ex_ d4 and OT-I T_ex_ d14 T cells as determined by quantitative RT-PCR and corrected for *Gapdh.* All data are mean ± SEM. C) ATACseq signal tracks at the *Pdcd1*, *Cd38, Havcr2, Ctla4, Lag3* and *Entpd1* loci highlighting differential chromatin accessibility peaks in T_ex_ d14 CD8^+^ T cells compared to splenic CD44^−^ T_naïve_ CD8^+^ cells.

**Figure S3.**
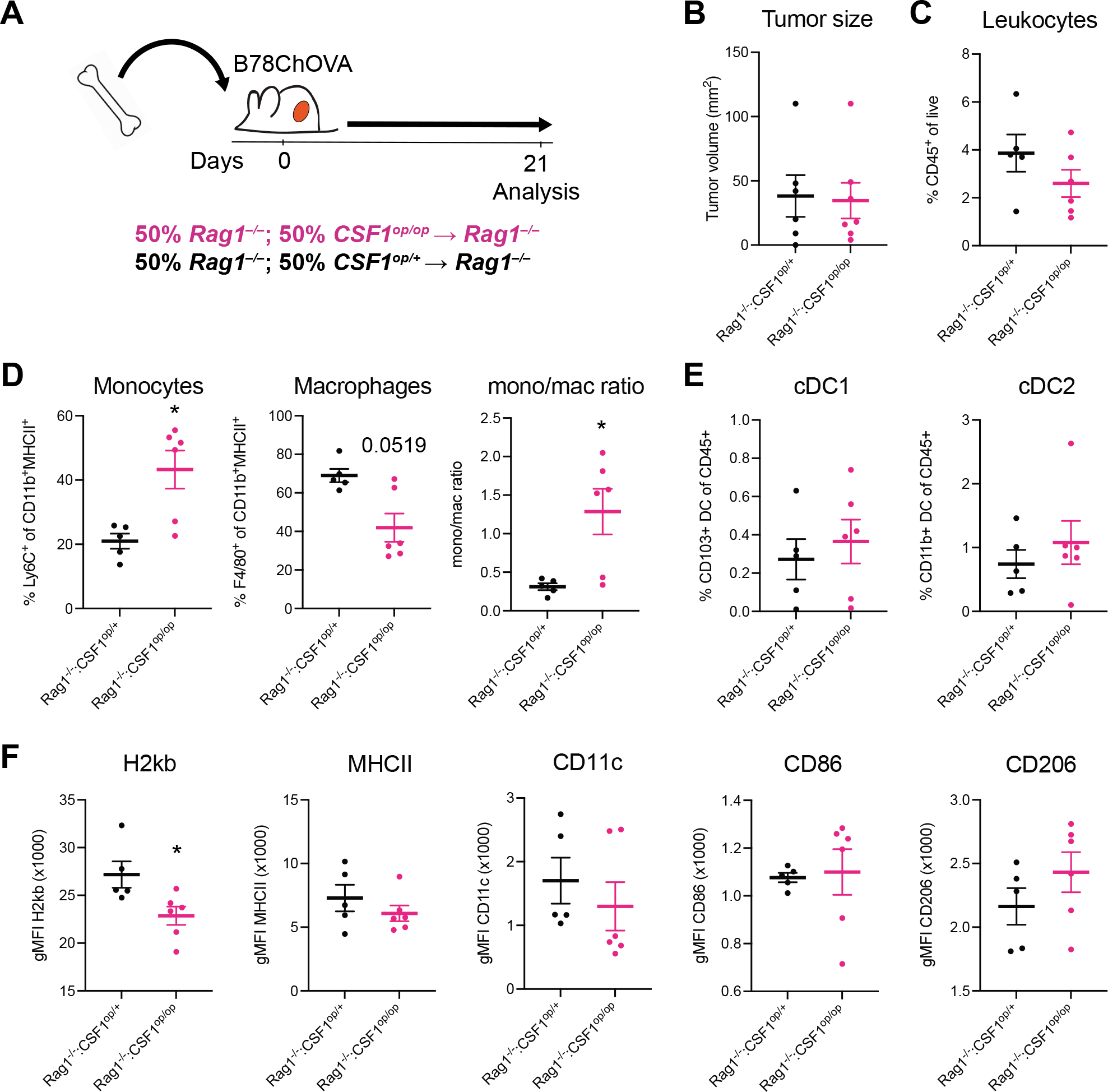
T cell-derived CSF1 shapes monocyte-macrophage dynamics in the TME (related to Figure 3). A) Experimental set-up of mixed bone marrow chimeras, reconstituted with a 50:50 mixture of *Rag1^−/∓^: CSF1^op/+^* (n = 5 mice) or *Rag1^−/−^: CSF1^op/op^* (n = 6 mice) inoculated with subcutaneous B78ChOVA melanomas 6-10 weeks after bone marrow reconstitution. 21 days later, mice were sacrificed for analysis of immune composition of tumors. B) Quantification of tumor volume (mm^2^) by caliper measurements at time of sacrifice. C-E) Flow cytometric analysis of total tumor-infiltrating CD45^+^ leukocytes (C) and (D) the proportion of Ly6C^+^ monocytes (left), F4/80^+^ macrophages (middle) of CD11b^+^MHCII^+^ cells and monocyte/macrophage ratio (right). E) Proportion of CD103^+^ cDC1 and CD11b^+^ cDC2 of total CD45^+^ cells. F) Quantification of expression of H2Kb, MHCII, CD11c, CD86 and CD206 gated on CD11b^+^ F4/80^+^ macrophages (gMFI) in B78ChOVA melanomas. Representative of two independent experiments. All data are mean ± SEM. Statistical significance was determined using the Mann-Whitney U test. * p < 0.05.

**Figure S4.**
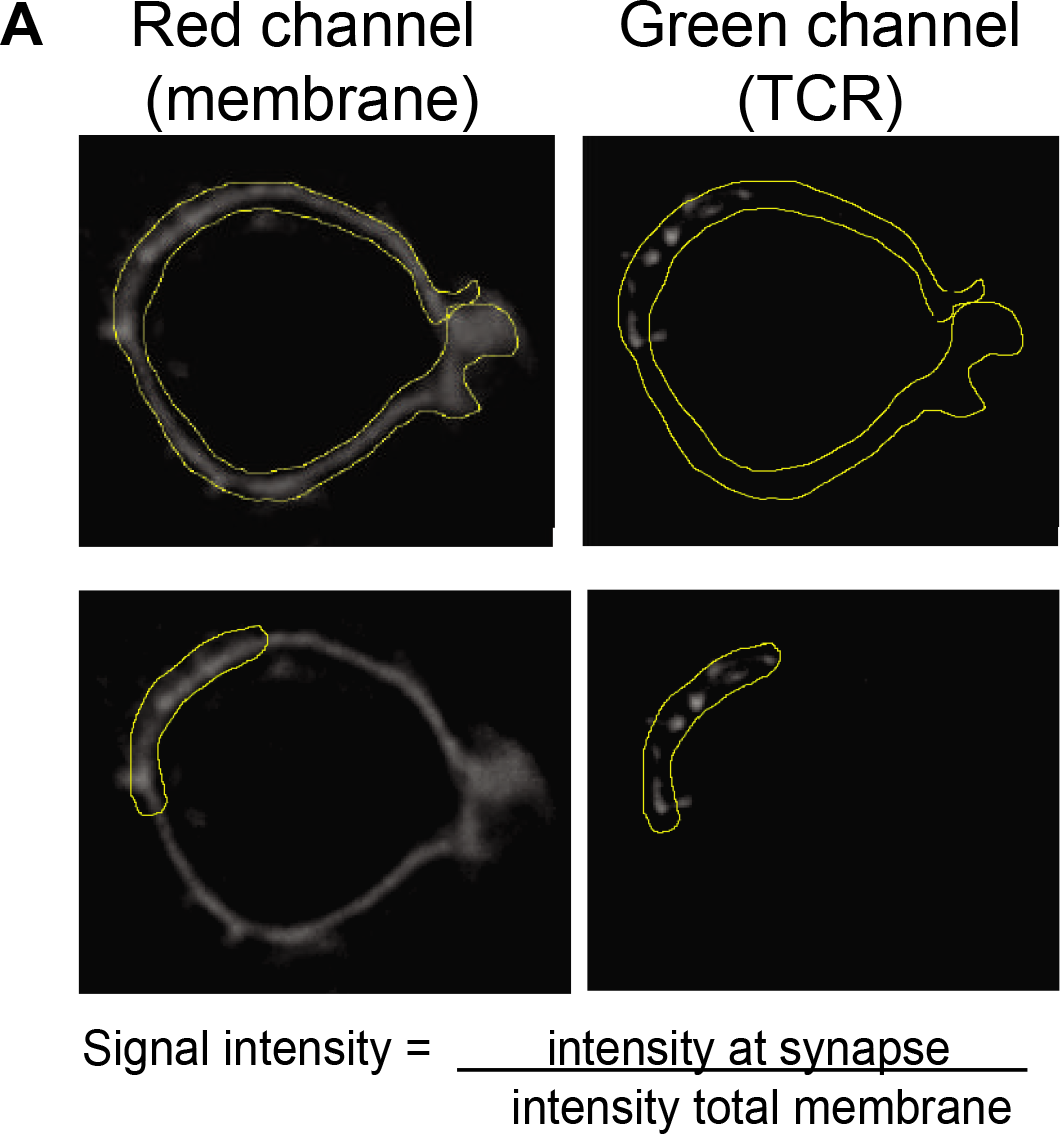
Synaptic TAM-CD8^+^ T cell interactions induce TCR clustering (related to Figure 4). A) TCR clustering on the T cell membrane was quantified by manually outlining the total T cell membrane versus TAM interaction site (synapse). Signal intensity for red (membrane) and green (TCR) channels were determined using ImageJ, and the ratio of signal intensity (synapse/total membrane) was calculated.

**Figure S5.**
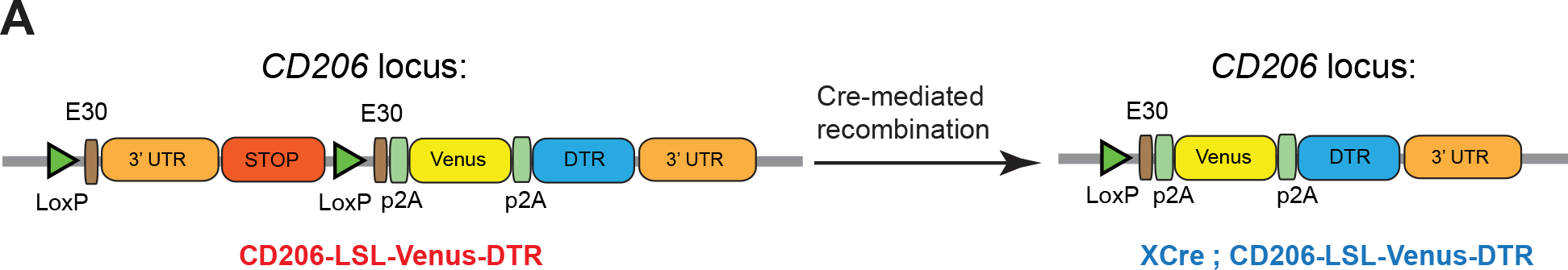
Schematic representation of a novel CD206-LSL-Venus-DTR reporter mouse model (related to Figure 6). Schematic representation of the genetic constructs used to generate a novel CD206-LSL-Venus-DTR reporter mouse model. Strain was crossed to CSF1R^Cre^ background to establish conditional deletion of the LSL cassette resulting in CD206-Venus expression specifically in myeloid cells.

**Figure S6.**
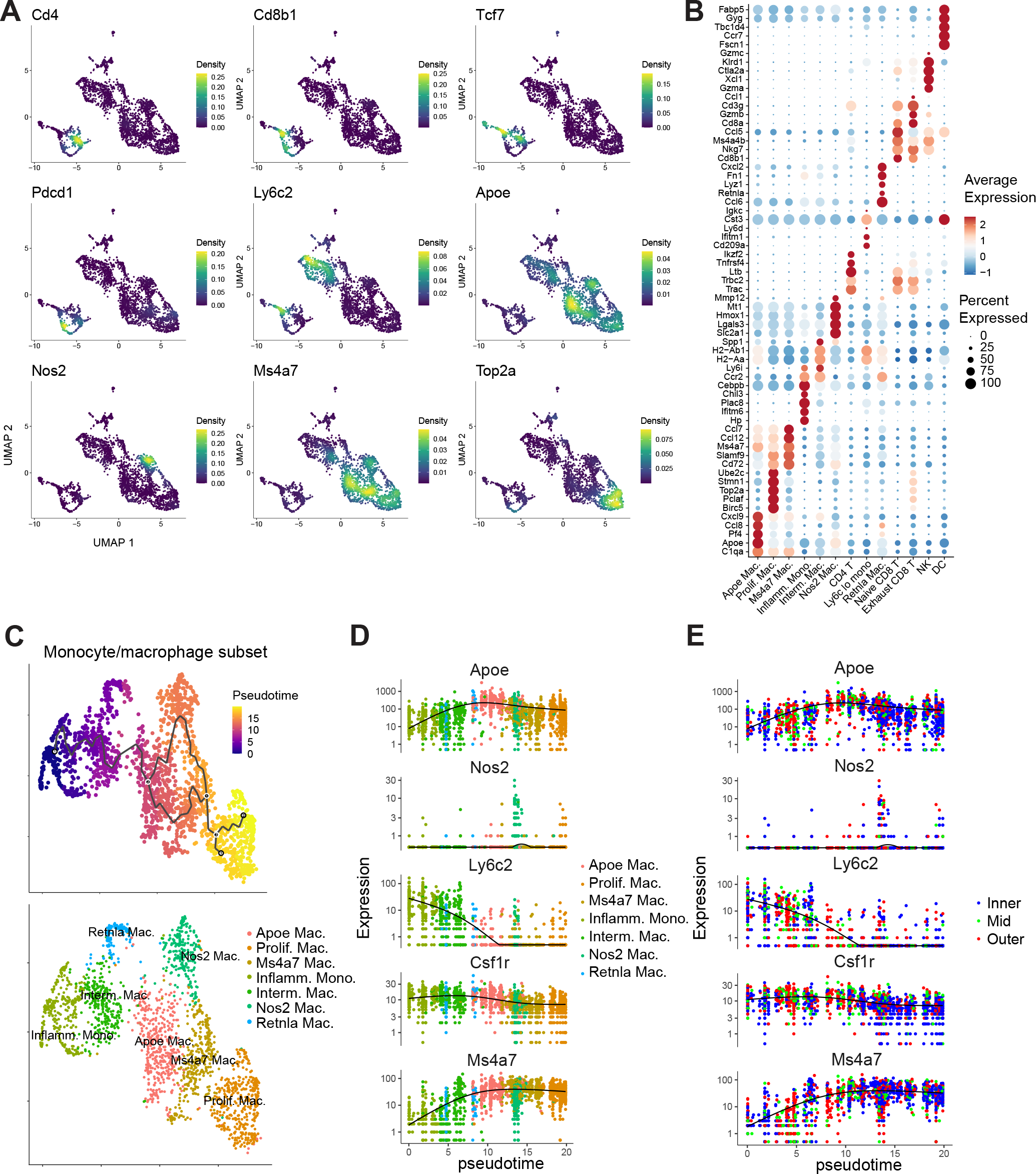
ZipSeq to spatially delineate TAM-T_ex_ interactions in the TME (related to Figure 6). A) Feature plots for selected marker genes using kernel density estimates (implemented by package ‘Nebulosa’ (Alquicira-Hernandez and Powell, 2021)) with *Cd4* (marking CD4^+^ T cells), *Cd8b1* (CD8^+^ T cells), *Tcf7* (naïve CD8^+^ T cells), *Pdcd1* (exhausted CD8^+^ T cells), *Ly6c2* (monocytes), *Apoe, Nos2, Ms4a7* and *Top2* (distinct macrophage subsets). B) Dotplot representation of marker gene expression (top 5 differentially expressed genes by LogFC expressed in at least 10% of cells) in annotated clusters. Dot size represents percent expression in cluster and color indicates average expression level. C) UMAP representation of monocyte/macrophage population state identity (lower) overlaid with pseudotime false-color through Monocle (upper), with *Ly6c2^HI^* inflammatory monocyte state designated as the root state. D) Expression of genes marking distinct monocyte/macrophage populations with increasing pseudotime demonstrating that cells within our defined trajectory lose expression of *Ly6c2* while gaining expression of *Apoe* and *Ms4a7* while maintaining *Csf1r* expression. E) Pseudotime plots from D overlaid with regional localization of monocyte/macrophage subsets in B78ChOVA tumors demonstrating that *Ly6c2^HI^* inflammatory monocytes are predominantly localized in the outer regions, while *Apoe^HI^* and *Ms4a7^HI^* macrophages are highly enriched in the inner regions of the TME (n = 2083 cells).

---

**Table S1.**
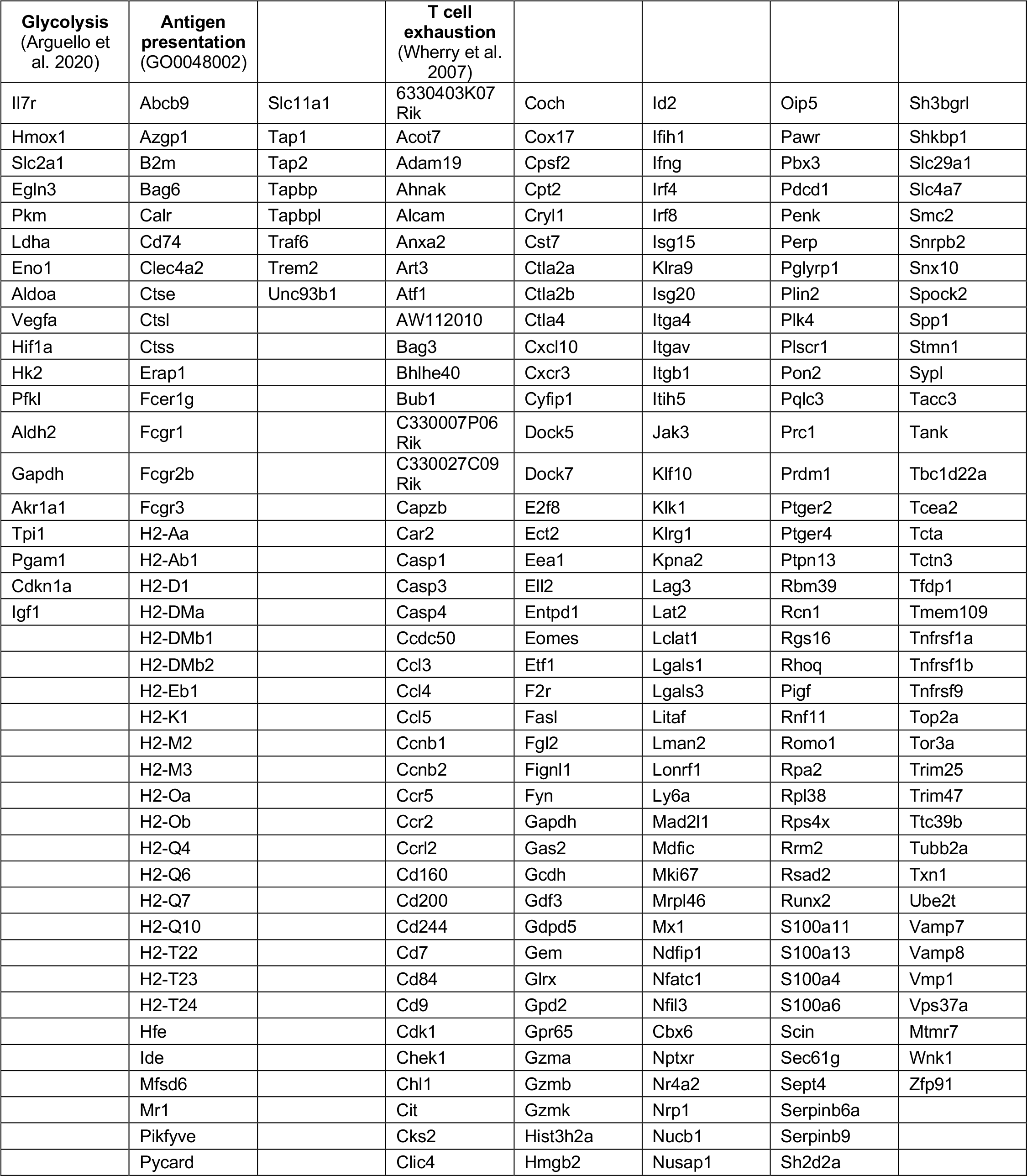
Gene lists used in ZipSeq related to Figure 6 and Figure S6.

## Key Resource Table

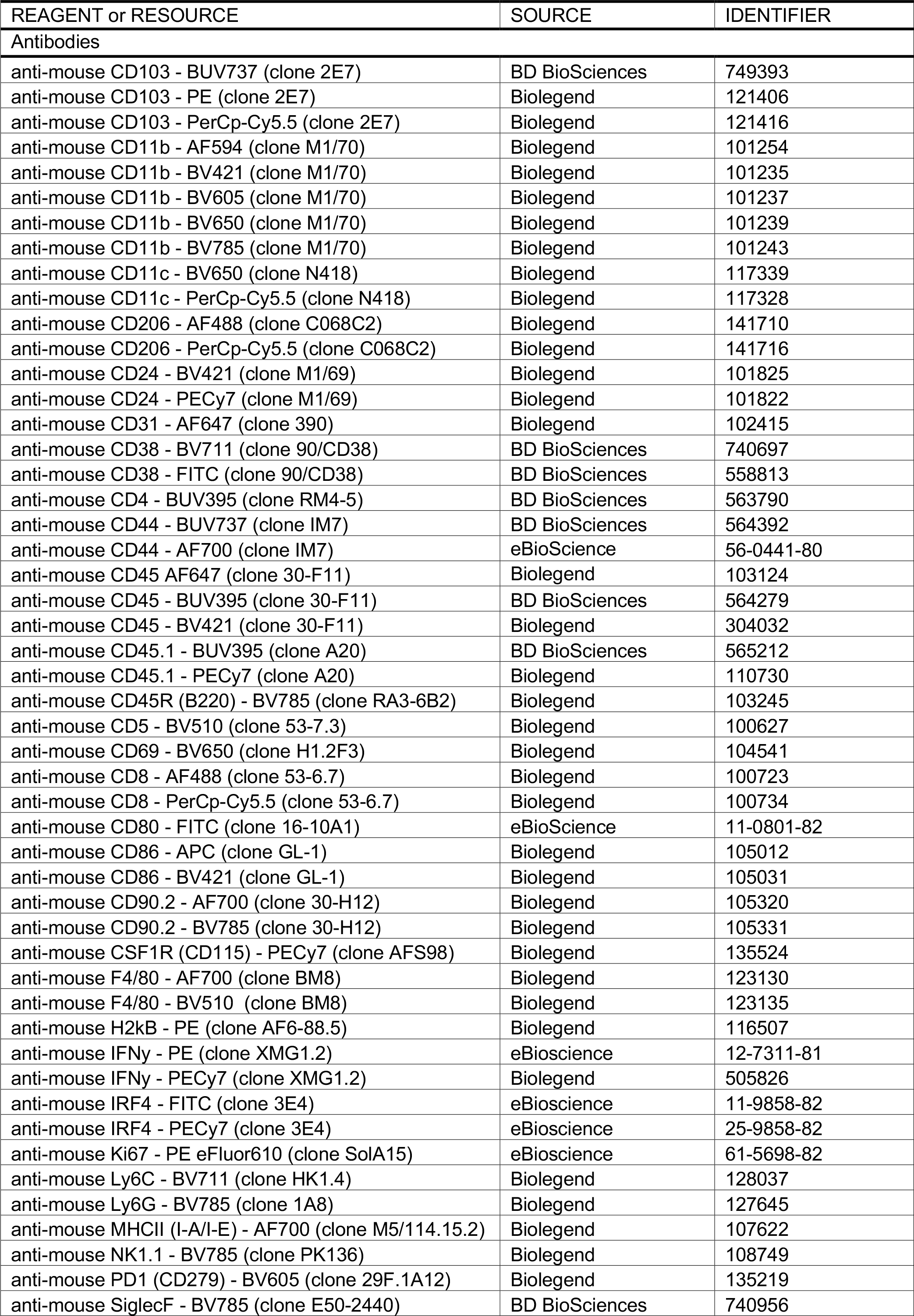
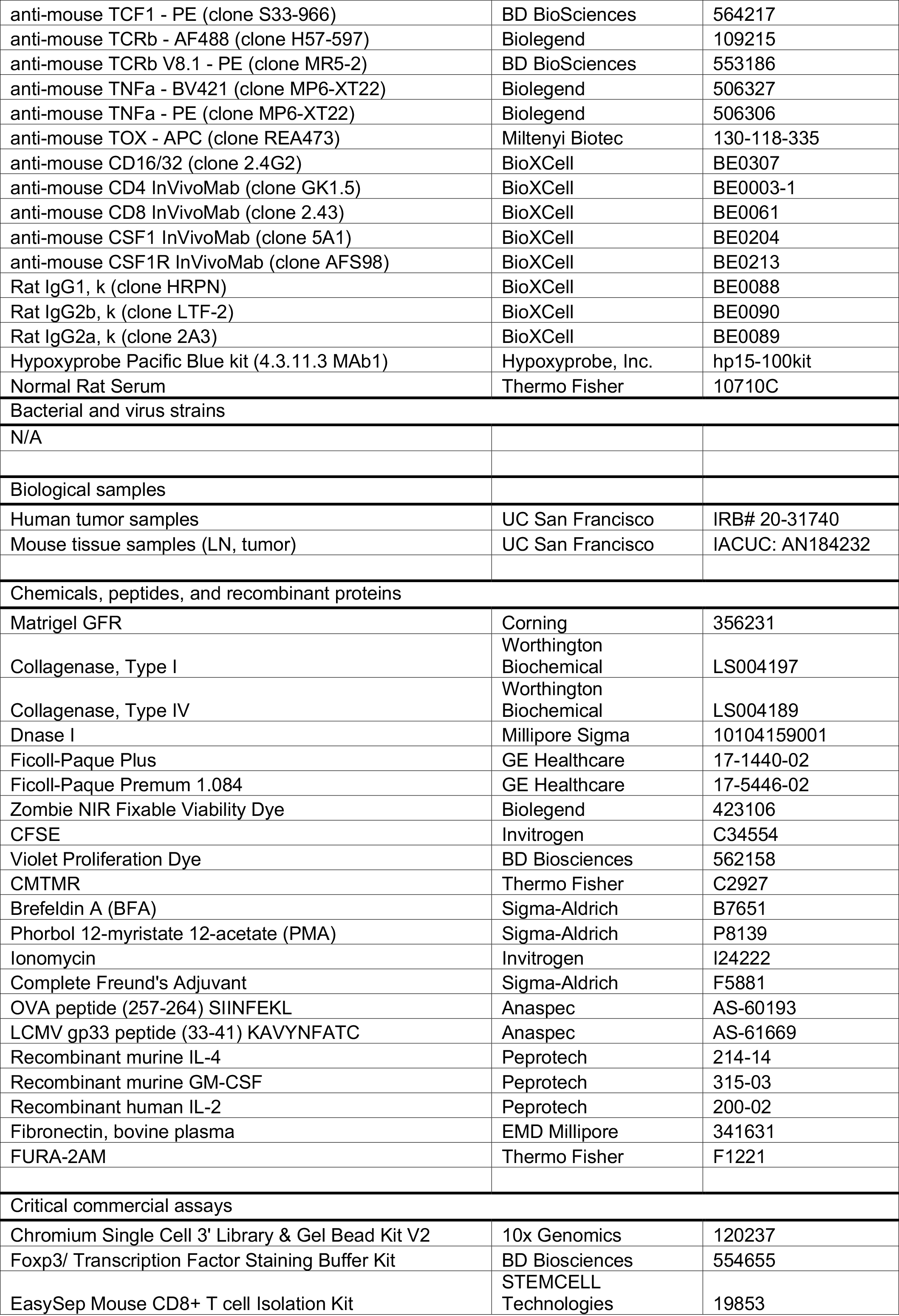
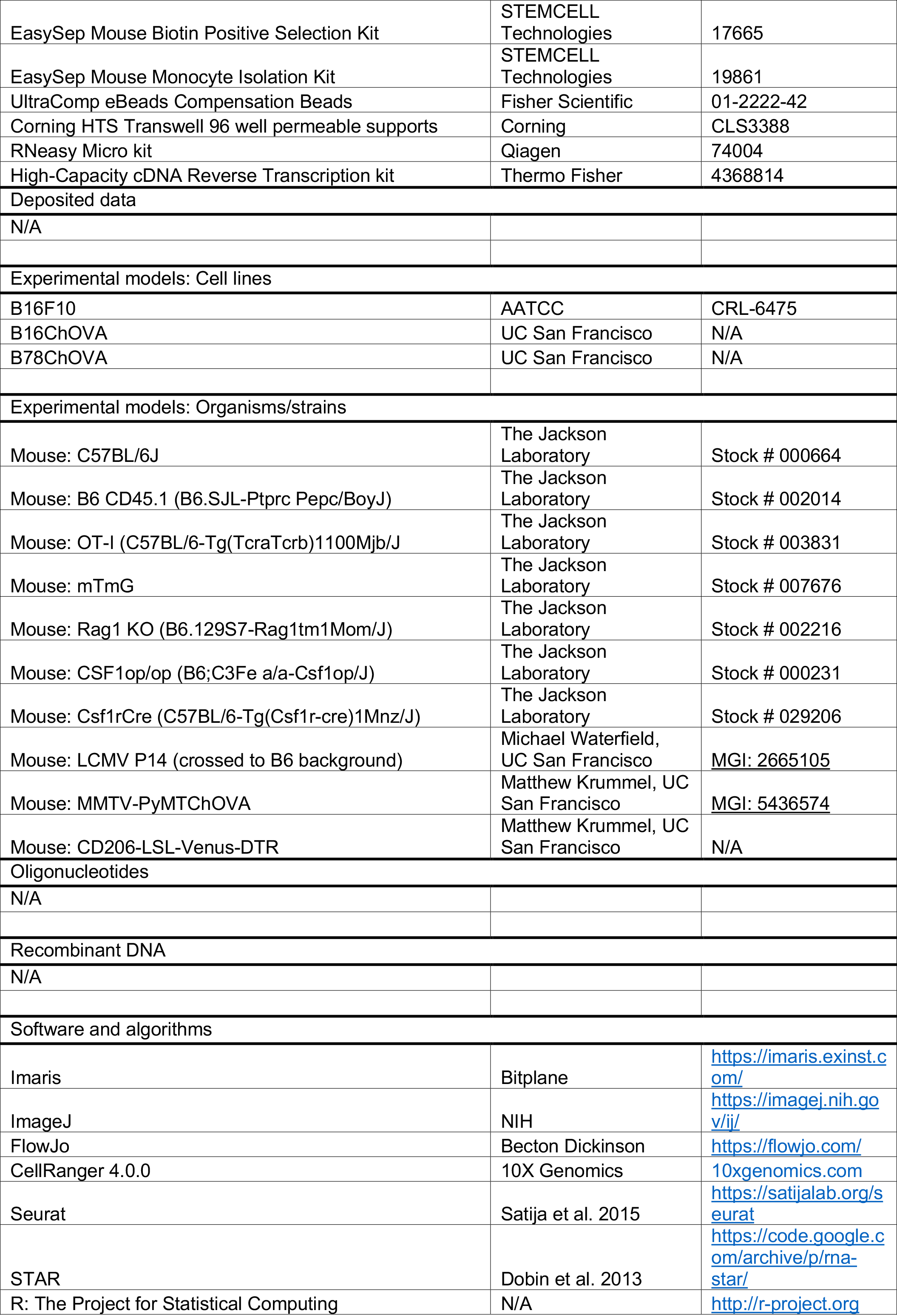
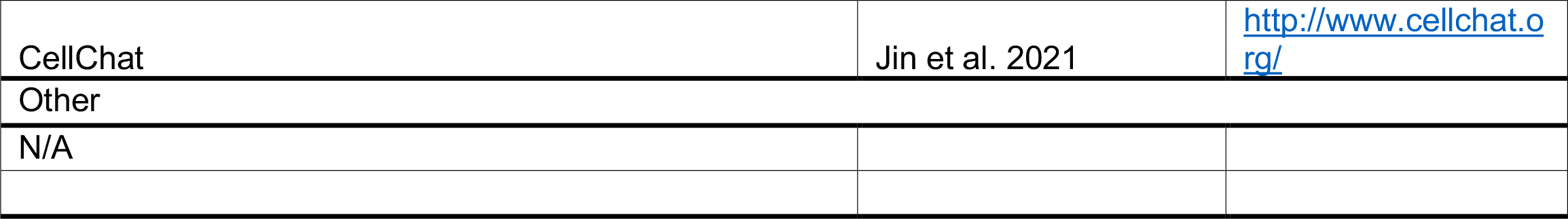

## References

Alfei, F., Kanev, K., Hofmann, M., Wu, M., Ghoneim, H.E., Roelli, P., Utzschneider, D.T., von Hoesslin, M., Cullen, J.G., Fan, Y., et al. (2019). TOX reinforces the phenotype and longevity of exhausted T cells in chronic viral infection. Nature 571, 265–269.

Alquicira-Hernandez, J., and Powell, J.E. (2021). Nebulosa recovers single cell gene expression signals by kernel density estimation. Bioinforma. Oxf. Engl. btab003.

Argüello, R.J., Combes, A.J., Char, R., Gigan, J.-P., Baaziz, A.I., Bousiquot, E., Camosseto, V., Samad, B., Tsui, J., Yan, P., et al. (2020). SCENITH: A Flow Cytometry-Based Method to Functionally Profile Energy Metabolism with Single-Cell Resolution. Cell Metab. 32, 1063–1075.

Barry, K.C., Hsu, J., Broz, M.L., Cueto, F.J., Binnewies, M., Combes, A.J., Nelson, A.E., Loo, K., Kumar, R., Rosenblum, M.D., et al. (2018). A natural killer-dendritic cell axis defines checkpoint therapy-responsive tumor microenvironments. Nat. Med. 24, 1178–1191.

Beatty, G.L., Chiorean, E.G., Fishman, M.P., Saboury, B., Teitelbaum, U.R., Sun, W., Huhn, R.D., Song, W., Li, D., Sharp, L.L., et al. (2011). CD40 agonists alter tumor stroma and show efficacy against pancreatic carcinoma in mice and humans. Science 331, 1612–1616.

Beltra, J.-C., Manne, S., Abdel-Hakeem, M.S., Kurachi, M., Giles, J.R., Chen, Z., Casella, V., Ngiow, S.F., Khan, O., Huang, Y.J., et al. (2020). Developmental Relationships of Four Exhausted CD8+ T Cell Subsets Reveals Underlying Transcriptional and Epigenetic Landscape Control Mechanisms. Immunity 52, 825–841.

Bengsch, B., Johnson, A.L., Kurachi, M., Odorizzi, P.M., Pauken, K.E., Attanasio, J., Stelekati, E., McLane, L.M., Paley, M.A., Delgoffe, G.M., et al. (2016). Bioenergetic Insufficiencies Due to Metabolic Alterations Regulated by the Inhibitory Receptor PD-1 Are an Early Driver of CD8(+) T Cell Exhaustion. Immunity 45, 358–373.

Bi, K., He, M.X., Bakouny, Z., Kanodia, A., Napolitano, S., Wu, J., Grimaldi, G., Braun, D.A., Cuoco, M.S., Mayorga, A., et al. (2021). Tumor and immune reprogramming during immunotherapy in advanced renal cell carcinoma. Cancer Cell 39, 649–661.

Binnewies, M., Roberts, E.W., Kersten, K., Chan, V., Fearon, D.F., Merad, M., Coussens, L.M., Gabrilovich, D.I., Ostrand-Rosenberg, S., Hedrick, C.C., et al. (2018). Understanding the tumor immune microenvironment (TIME) for effective therapy. Nat. Med. 24, 541–550.

Binnewies, M., Mujal, A.M., Pollack, J.L., Combes, A.J., Hardison, E.A., Barry, K.C., Tsui, J., Ruhland, M.K., Kersten, K., Abushawish, M.A., et al. (2019). Unleashing Type-2 Dendritic Cells to Drive Protective Antitumor CD4+ T Cell Immunity. Cell 177, 556–571.

Boissonnas, A., Licata, F., Poupel, L., Jacquelin, S., Fetler, L., Krumeich, S., Théry, C., Amigorena, S., and Combadière, C. (2013). CD8+ tumor-infiltrating T cells are trapped in the tumor-dendritic cell network. Neoplasia N. Y. N 15, 85–94.

Boldajipour, B., Nelson, A., and Krummel, M.F. (2016). Tumor-infiltrating lymphocytes are dynamically desensitized to antigen but are maintained by homeostatic cytokine. JCI Insight 1, e89289.

Braun, D.A., Street, K., Burke, K.P., Cookmeyer, D.L., Denize, T., Pedersen, C.B., Gohil, S.H., Schindler, N., Pomerance, L., Hirsch, L., et al. (2021). Progressive immune dysfunction with advancing disease stage in renal cell carcinoma. Cancer Cell 39, 632–648.

Broz, M.L., Binnewies, M., Boldajipour, B., Nelson, A.E., Pollack, J.L., Erle, D.J., Barczak, A., Rosenblum, M.D., Daud, A., Barber, D.L., et al. (2014). Dissecting the Tumor Myeloid Compartment Reveals Rare Activating Antigen-Presenting Cells Critical for T Cell Immunity. Cancer Cell 26, 638–652.

Buechler, M.B., Fu, W., and Turley, S.J. (2021). Fibroblast-macrophage reciprocal interactions in health, fibrosis, and cancer. Immunity 54, 903–915.

Cai, E., Marchuk, K., Beemiller, P., Beppler, C., Rubashkin, M.G., Weaver, V.M., Gérard, A., Liu, T.-L., Chen, B.-C., Betzig, E., et al. (2017). Visualizing dynamic microvillar search and stabilization during ligand detection by T cells. Science 356, eaal3118.

Cao, J., Spielmann, M., Qiu, X., Huang, X., Ibrahim, D.M., Hill, A.J., Zhang, F., Mundlos, S., Christiansen, L., Steemers, F.J., et al. (2019). The single-cell transcriptional landscape of mammalian organogenesis. Nature 566, 496–502.

Chen, S., Zhou, Y., Chen, Y., and Gu, J. (2018). fastp: an ultra-fast all-in-one FASTQ preprocessor. Bioinformatics 34, i884– i890.

Chen, Y., Zander, R.A., Wu, X., Schauder, D.M., Kasmani, M.Y., Shen, J., Zheng, S., Burns, R., Taparowsky, E.J., and Cui, W. (2021). BATF regulates progenitor to cytolytic effector CD8+ T cell transition during chronic viral infection. Nat. Immunol. 22, 996–1007.

Cheng, S., Li, Z., Gao, R., Xing, B., Gao, Y., Yang, Y., Qin, S., Zhang, L., Ouyang, H., Du, P., et al. (2021). A pan-cancer single-cell transcriptional atlas of tumor infiltrating myeloid cells. Cell 184, 792–809.

Combes, A.J., Samad, B., Tsui, J., Chew, N.W., Yan, P., Reeder, G.C., Kushnoor, D., Shen, A., Davidson, B., Barczac, A.J., et al. (2021). A Pan-Cancer Census of Dominant Tumor Immune Archetypes. BioRxiv 2021.04.26.441344.

Corces, M.R., Trevino, A.E., Hamilton, E.G., Greenside, P.G., Sinnott-Armstrong, N.A., Vesuna, S., Satpathy, A.T., Rubin, A.J., Montine, K.S., Wu, B., et al. (2017). An improved ATAC-seq protocol reduces background and enables interrogation of frozen tissues. Nat. Methods 14, 959–962.

Corces, M.R., Granja, J.M., Shams, S., Louie, B.H., Seoane, J.A., Zhou, W., Silva, T.C., Groeneveld, C., Wong, C.K., Cho, S.W., et al. (2018). The chromatin accessibility landscape of primary human cancers. Science 362, eaav1898.

De Palma, M., and Lewis, C.E. (2013). Macrophage regulation of tumor responses to anticancer therapies. Cancer Cell 23, 277–286.

DeNardo, D.G., and Ruffell, B. (2019). Macrophages as regulators of tumour immunity and immunotherapy. Nat. Rev. Immunol. 19, 369–382.

DeNardo, D.G., Brennan, D.J., Rexhepaj, E., Ruffell, B., Shiao, S.L., Madden, S.F., Gallagher, W.M., Wadhwani, N., Keil, S.D., Junaid, S.A., et al. (2011). Leukocyte Complexity Predicts Breast Cancer Survival and Functionally Regulates Response to Chemotherapy. Cancer Discov. 1, 54–67.

Dobin, A., Davis, C.A., Schlesinger, F., Drenkow, J., Zaleski, C., Jha, S., Batut, P., Chaisson, M., and Gingeras, T.R. (2013). STAR: ultrafast universal RNA-seq aligner. Bioinformatics 29, 15–21.

Doering, T.A., Crawford, A., Angelosanto, J.M., Paley, M.A., Ziegler, C.G., and Wherry, E.J. (2012). Network analysis reveals centrally connected genes and pathways involved in CD8+ T cell exhaustion versus memory. Immunity 37, 1130– 1144.

Engelhardt, J.J., Boldajipour, B., Beemiller, P., Pandurangi, P., Sorensen, C., Werb, Z., Egeblad, M., and Krummel, M.F. (2012). Marginating dendritic cells of the tumor microenvironment cross-present tumor antigens and stably engage tumor-specific T cells. Cancer Cell 21, 402–417.

Frankish, A., Diekhans, M., Ferreira, A.-M., Johnson, R., Jungreis, I., Loveland, J., Mudge, J.M., Sisu, C., Wright, J., Armstrong, J., et al. (2019). GENCODE reference annotation for the human and mouse genomes. Nucleic Acids Res. 47, D766–D773.

Galon, J., Costes, A., Sanchez-Cabo, F., Kirilovsky, A., Mlecnik, B., Lagorce-Pagès, C., Tosolini, M., Camus, M., Berger, A., Wind, P., et al. (2006). Type, density, and location of immune cells within human colorectal tumors predict clinical outcome. Science 313, 1960–1964.

Gentles, A.J., Newman, A.M., Liu, C.L., Bratman, S.V., Feng, W., Kim, D., Nair, V.S., Xu, Y., Khuong, A., Hoang, C.D., et al. (2015). The prognostic landscape of genes and infiltrating immune cells across human cancers. Nat. Med. 21, 938–945.

Guerriero, J.L. (2019). Macrophages: Their Untold Story in T Cell Activation and Function. Int. Rev. Cell Mol. Biol. 342, 73– 93.

Hafemeister, C., and Satija, R. (2019). Normalization and variance stabilization of single-cell RNA-seq data using regularized negative binomial regression. Genome Biol. 20, 296.

Hong, F., Meng, Q., Zhang, W., Zheng, R., Li, X., Cheng, T., Hu, D., and Gao, X. (2021). Single-Cell Analysis of the Pan-Cancer Immune Microenvironment and scTIME Portal. Cancer Immunol. Res. 9, 939–951.

Hu, K.H., Eichorst, J.P., McGinnis, C.S., Patterson, D.M., Chow, E.D., Kersten, K., Jameson, S.C., Gartner, Z.J., Rao, A.A., and Krummel, M.F. (2020). ZipSeq: barcoding for real-time mapping of single cell transcriptomes. Nat. Methods 17, 833– 843.

Im, S.J., Hashimoto, M., Gerner, M.Y., Lee, J., Kissick, H.T., Burger, M.C., Shan, Q., Hale, J.S., Lee, J., Nasti, T.H., et al. (2016). Defining CD8+ T cells that provide the proliferative burst after PD-1 therapy. Nature 537, 417–421.

Jansen, C.S., Prokhnevska, N., Master, V.A., Sanda, M.G., Carlisle, J.W., Bilen, M.A., Cardenas, M., Wilkinson, S., Lake, R., Sowalsky, A.G., et al. (2019). An intra-tumoral niche maintains and differentiates stem-like CD8 T cells. Nature 576, 465–470.

Jin, S., Guerrero-Juarez, C.F., Zhang, L., Chang, I., Ramos, R., Kuan, C.-H., Myung, P., Plikus, M.V., and Nie, Q. (2021). Inference and analysis of cell-cell communication using CellChat. Nat. Commun. 12, 1088.

Katzenelenbogen, Y., Sheban, F., Yalin, A., Yofe, I., Svetlichnyy, D., Jaitin, D.A., Bornstein, C., Moshe, A., Keren-Shaul, H., Cohen, M., et al. (2020). Coupled scRNA-Seq and Intracellular Protein Activity Reveal an Immunosuppressive Role of TREM2 in Cancer. Cell 182, 872–885.

Khan, O., Giles, J.R., McDonald, S., Manne, S., Ngiow, S.F., Patel, K.P., Werner, M.T., Huang, A.C., Alexander, K.A., Wu, J.E., et al. (2019). TOX transcriptionally and epigenetically programs CD8 + T cell exhaustion. Nature 571, 211.

Klug, F., Prakash, H., Huber, P.E., Seibel, T., Bender, N., Halama, N., Pfirschke, C., Voss, R.H., Timke, C., Umansky, L., et al. (2013). Low-dose irradiation programs macrophage differentiation to an iNOS^+^/M1 phenotype that orchestrates effective T cell immunotherapy. Cancer Cell 24, 589–602.

Law, C.W., Chen, Y., Shi, W., and Smyth, G.K. (2014). voom: precision weights unlock linear model analysis tools for RNA-seq read counts. Genome Biol. 15, R29.

Man, K., Gabriel, S.S., Liao, Y., Gloury, R., Preston, S., Henstridge, D.C., Pellegrini, M., Zehn, D., Berberich-Siebelt, F., Febbraio, M.A., et al. (2017). Transcription Factor IRF4 Promotes CD8+ T Cell Exhaustion and Limits the Development of Memory-like T Cells during Chronic Infection. Immunity 47, 1129–1141.

Miller, B.C., Sen, D.R., Al Abosy, R., Bi, K., Virkud, Y.V., LaFleur, M.W., Yates, K.B., Lako, A., Felt, K., Naik, G.S., et al. (2019). Subsets of exhausted CD8+ T cells differentially mediate tumor control and respond to checkpoint blockade. Nat. Immunol. 20, 326–336.

Molgora, M., Esaulova, E., Vermi, W., Hou, J., Chen, Y., Luo, J., Brioschi, S., Bugatti, M., Omodei, A.S., Ricci, B., et al. (2020). TREM2 Modulation Remodels the Tumor Myeloid Landscape Enhancing Anti-PD-1 Immunotherapy. Cell 182, 886– 900.

Mujal, A.M., Combes, A.J., Rao, A.R., Binnewies, M., Samad, B., Tsui, J., Boissonnas, A., Pollack, J.L., Argüello, R.J., Ruhland, M.K., et al. (2021). Holistic Characterization of Tumor Monocyte-to-Macrophage Differentiation Integrates Distinct Immune Phenotypes in Kidney Cancer. BioRxiv 2021.07.07.451502.

O’Connell, P., Hyslop, S., Blake, M.K., Godbehere, S., Amalfitano, A., and Aldhamen, Y.A. (2021). SLAMF7 Signaling Reprograms T Cells toward Exhaustion in the Tumor Microenvironment. J. Immunol. Baltim. Md 1950 *206*, 193–205.

Oliveira, G., Stromhaug, K., Klaeger, S., Kula, T., Frederick, D.T., Le, P.M., Forman, J., Huang, T., Li, S., Zhang, W., et al. (2021). Phenotype, specificity and avidity of antitumour CD8+ T cells in melanoma. Nature 596, 119–125.

Pauken, K.E., Sammons, M.A., Odorizzi, P.M., Manne, S., Godec, J., Khan, O., Drake, A.M., Chen, Z., Sen, D.R., Kurachi, M., et al. (2016). Epigenetic stability of exhausted T cells limits durability of reinvigoration by PD-1 blockade. Science 354, 1160–1165.

Peranzoni, E., Lemoine, J., Vimeux, L., Feuillet, V., Barrin, S., Kantari-Mimoun, C., Bercovici, N., Guérin, M., Biton, J., Ouakrim, H., et al. (2018). Macrophages impede CD8 T cells from reaching tumor cells and limit the efficacy of anti–PD-1 treatment. Proc. Natl. Acad. Sci. 115, E4041–E4050.

Philip, M., and Schietinger, A. (2021). CD8+ T cell differentiation and dysfunction in cancer. Nat. Rev. Immunol.

Philip, M., Fairchild, L., Sun, L., Horste, E.L., Camara, S., Shakiba, M., Scott, A.C., Viale, A., Lauer, P., Merghoub, T., et al. (2017). Chromatin states define tumour-specific T cell dysfunction and reprogramming. Nature 545, 452–456.

Pritykin, Y., van der Veeken, J., Pine, A.R., Zhong, Y., Sahin, M., Mazutis, L., Pe’er, D., Rudensky, A.Y., and Leslie, C.S. (2021). A unified atlas of CD8 T cell dysfunctional states in cancer and infection. Mol. Cell 81, 2477–2493.

Roberts, E.W., Broz, M.L., Binnewies, M., Headley, M.B., Nelson, A.E., Wolf, D.M., Kaisho, T., Bogunovic, D., Bhardwaj, N., and Krummel, M.F. (2016). Critical Role for CD103+/CD141+ Dendritic Cells Bearing CCR7 for Tumor Antigen Trafficking and Priming of T Cell Immunity in Melanoma. Cancer Cell 30, 324–336.

Robinson, M.D., McCarthy, D.J., and Smyth, G.K. (2010). edgeR: a Bioconductor package for differential expression analysis of digital gene expression data. Bioinformatics 26, 139–140.

Ruffell, B., Au, A., Rugo, H.S., Esserman, L.J., Hwang, E.S., and Coussens, L.M. (2012). Leukocyte composition of human breast cancer. Proc. Natl. Acad. Sci. U. S. A. 109, 2796–2801.

Sade-Feldman, M., Yizhak, K., Bjorgaard, S.L., Ray, J.P., de Boer, C.G., Jenkins, R.W., Lieb, D.J., Chen, J.H., Frederick, D.T., Barzily-Rokni, M., et al. (2018). Defining T Cell States Associated with Response to Checkpoint Immunotherapy in Melanoma. Cell 175, 998–1013.

Salmon, H., Idoyaga, J., Rahman, A., Leboeuf, M., Remark, R., Jordan, S., Casanova-Acebes, M., Khudoynazarova, M., Agudo, J., Tung, N., et al. (2016). Expansion and Activation of CD103+ Dendritic Cell Progenitors at the Tumor Site Enhances Tumor Responses to Therapeutic PD-L1 and BRAF Inhibition. Immunity 44, 924–938.

Satija, R., Farrell, J.A., Gennert, D., Schier, A.F., and Regev, A. (2015). Spatial reconstruction of single-cell gene expression data. Nat. Biotechnol. 33, 495–502.

Satpathy, A.T., Granja, J.M., Yost, K.E., Qi, Y., Meschi, F., McDermott, G.P., Olsen, B.N., Mumbach, M.R., Pierce, S.E., Corces, M.R., et al. (2019). Massively parallel single-cell chromatin landscapes of human immune cell development and intratumoral T cell exhaustion. Nat. Biotechnol. 37, 925–936.

Scharping, N.E., Rivadeneira, D.B., Menk, A.V., Vignali, P.D.A., Ford, B.R., Rittenhouse, N.L., Peralta, R., Wang, Y., Wang, Y., DePeaux, K., et al. (2021). Mitochondrial stress induced by continuous stimulation under hypoxia rapidly drives T cell exhaustion. Nat. Immunol. 22, 205–215.

Schietinger, A., Philip, M., Krisnawan, V.E., Chiu, E.Y., Delrow, J.J., Basom, R.S., Lauer, P., Brockstedt, D.G., Knoblaugh, S.E., Hämmerling, G.J., et al. (2016). Tumor-Specific T Cell Dysfunction Is a Dynamic Antigen-Driven Differentiation Program Initiated Early during Tumorigenesis. Immunity 45, 389–401.

Scott, A.C., Dündar, F., Zumbo, P., Chandran, S.S., Klebanoff, C.A., Shakiba, M., Trivedi, P., Menocal, L., Appleby, H., Camara, S., et al. (2019). TOX is a critical regulator of tumour-specific T cell differentiation. Nature 571, 270–274.

Seo, H., González-Avalos, E., Zhang, W., Ramchandani, P., Yang, C., Lio, C.-W.J., Rao, A., and Hogan, P.G. (2021). BATF and IRF4 cooperate to counter exhaustion in tumor-infiltrating CAR T cells. Nat. Immunol. 22, 983–995.

Siddiqui, I., Schaeuble, K., Chennupati, V., Fuertes Marraco, S.A., Calderon-Copete, S., Pais Ferreira, D., Carmona, S.J., Scarpellino, L., Gfeller, D., Pradervand, S., et al. (2019). Intratumoral Tcf1+PD-1+CD8+ T Cells with Stem-like Properties Promote Tumor Control in Response to Vaccination and Checkpoint Blockade Immunotherapy. Immunity 50, 195–211.

Smyth, G.K. (2005). limma: Linear Models for Microarray Data. In Bioinformatics and Computational Biology Solutions Using R and Bioconductor, R. Gentleman, V.J. Carey, W. Huber, R.A. Irizarry, and S. Dudoit, eds. (New York, NY: Springer New York), pp. 397–420.

Spranger, S., Dai, D., Horton, B., and Gajewski, T.F. (2017). Tumor-Residing Batf3 Dendritic Cells Are Required for Effector T Cell Trafficking and Adoptive T Cell Therapy. Cancer Cell 31, 711–723.

Thommen, D.S., Koelzer, V.H., Herzig, P., Roller, A., Trefny, M., Dimeloe, S., Kiialainen, A., Hanhart, J., Schill, C., Hess, C., et al. (2018). A transcriptionally and functionally distinct PD-1+ CD8+ T cell pool with predictive potential in non-small-cell lung cancer treated with PD-1 blockade. Nat. Med. 24, 994–1004.

Tumeh, P.C., Harview, C.L., Yearley, J.H., Shintaku, I.P., Taylor, E.J.M., Robert, L., Chmielowski, B., Spasic, M., Henry, G., Ciobanu, V., et al. (2014). PD-1 blockade induces responses by inhibiting adaptive immune resistance. Nature 515, 568–571.

Utzschneider, D.T., Alfei, F., Roelli, P., Barras, D., Chennupati, V., Darbre, S., Delorenzi, M., Pinschewer, D.D., and Zehn, D. (2016). High antigen levels induce an exhausted phenotype in a chronic infection without impairing T cell expansion and survival. J. Exp. Med. 213, 1819–1834.

Utzschneider, D.T., Gabriel, S.S., Chisanga, D., Gloury, R., Gubser, P.M., Vasanthakumar, A., Shi, W., and Kallies, A. (2020). Early precursor T cells establish and propagate T cell exhaustion in chronic infection. Nat. Immunol. 21, 1256–1266.

Vardhana, S.A., Hwee, M.A., Berisa, M., Wells, D.K., Yost, K.E., King, B., Smith, M., Herrera, P.S., Chang, H.Y., Satpathy, A.T., et al. (2020). Impaired mitochondrial oxidative phosphorylation limits the self-renewal of T cells exposed to persistent antigen. Nat. Immunol. 21, 1022–1033.

Wagner, J., Rapsomaniki, M.A., Chevrier, S., Anzeneder, T., Langwieder, C., Dykgers, A., Rees, M., Ramaswamy, A., Muenst, S., Soysal, S.D., et al. (2019). A Single-Cell Atlas of the Tumor and Immune Ecosystem of Human Breast Cancer. Cell 177, 1330–1345.

Weber, E.W., Parker, K.R., Sotillo, E., Lynn, R.C., Anbunathan, H., Lattin, J., Good, Z., Belk, J.A., Daniel, B., Klysz, D., et al. (2021). Transient “rest” restores functionality in exhausted CAR-T cells via epigenetic remodeling. Science 372, eaba1786.

Wherry, E.J., Ha, S.-J., Kaech, S.M., Haining, W.N., Sarkar, S., Kalia, V., Subramaniam, S., Blattman, J.N., Barber, D.L., and Ahmed, R. (2007). Molecular signature of CD8+ T cell exhaustion during chronic viral infection. Immunity 27, 670–684.

Yao, C., Sun, H.-W., Lacey, N.E., Ji, Y., Moseman, E.A., Shih, H.-Y., Heuston, E.F., Kirby, M., Anderson, S., Cheng, J., et al. (2019). Single-cell RNA-seq reveals TOX as a key regulator of CD8+ T cell persistence in chronic infection. Nat. Immunol. 20, 890–901.

Zhang, Q., Liu, L., Gong, C., Shi, H., Zeng, Y., Wang, X., Zhao, Y., and Wei, Y. (2012). Prognostic significance of tumor-associated macrophages in solid tumor: a meta-analysis of the literature. PloS One 7, e50946.

